# Sensing, Feeling, and Regulating: Investigating the Association of Focal Brain Damage with Voluntary Respiratory and Motor Control

**DOI:** 10.1101/2023.10.16.562254

**Authors:** Henrik Bischoff, Christopher Kovach, Sukbhinder Kumar, Joel Bruss, Daniel Tranel, Sahib S Khalsa

**Affiliations:** Department of Psychology, University of Stockholm, Stockholm, Sweden; Department of Psychology, Carl-von-Ossietzky University Oldenburg, Oldenburg, Germany; Department of Neurosurgery, University of Iowa, Iowa City, USA; Department of Neurosurgery, University of Nebraska Medical Center, Omaha, NE, USA; Departments of Pediatrics, Neurology, and Psychiatry, University of Iowa, Iowa City, USA; Departments of Neurology and Psychological and Brain Sciences, University of Iowa, Iowa City, USA; Laureate Institute for Brain Research, Tulsa, OK, USA; Oxley College of Health Sciences, University of Tulsa, Tulsa, OK, USA

**Keywords:** respiration, interoception, brain damage, emotion, affect, amygdala, motor control

## Abstract

Breathing is a complex, vital function that can be modulated to influence physical and mental well-being. However, the role of cortical and subcortical brain regions in voluntary control of human respiration is underexplored. Here we investigated the influence of damage to human frontal, temporal, or limbic regions on the sensation and regulation of breathing patterns. Participants performed a respiratory regulation task across regular and irregular frequencies ranging from 6 to 60 breaths per minute (bpm), with a counterbalanced hand motor control task. Interoceptive and affective states induced by each condition were assessed via questionnaire and autonomic signals were indexed via skin conductance. Participants with focal lesions to the bilateral frontal lobe, right insula/basal ganglia, and left medial temporal lobe showed reduced performance than individually matched healthy comparisons during the breathing and motor tasks. They also reported significantly higher anxiety during the 60-bpm regular and irregular breathing trials, with anxiety correlating with difficulty in rapid breathing specifically within this group. This study demonstrates that damage to frontal, temporal, or limbic regions is associated with abnormal voluntary respiratory and motor regulation and tachypnea-related anxiety, highlighting the role of the forebrain in affective and motor responses during breathing.

**Highlights:**

- Impaired human respiratory regulation is associated with cortical/subcortical brain lesions
- Frontolimbic/temporal regions contribute to rhythmic breathing and hand motor control
- Frontolimbic/temporal damage is associated with anxiety during tachypnea/irregular breathing
- The human forebrain is vital for affective and interoceptive experiences during breathing

## Introduction

Breathing is a vital yet complex function performed continuously by vertebrates from birth until death. Crucial for cellular metabolism and gas exchange and often automatically performed, human breathing can be consciously controlled (Shweta, Pramod & Chen, 2021; Lumb, 2016), impacting physical (Hernandez, Manning & Zhang, 2019; Kim, Shin & Cho, 2022; Russo, Santarelli & O’Rourke, 2017) and mental health outcomes (Banushi et al., 2023; Balban et al., 2023) as well as central (Karjalainen, Kujala & Parviainen, 2023) and peripheral autonomic arousal (Laborde et al., 2022). Voluntary breathing refers to the conscious control of breathing, such as when we hold our breath, take a deep breath, or exhale forcefully. Voluntary breathing has been shown to modulate neural (McKay et al., 2003; Herrero et al., 2018), cognitive (Zaccaro et al., 2018) and emotional states (Jerath & Beveridge, 2020), demonstrating the relevance of this function to numerous health-related processes. While automatic breathing is presumptively generated by brainstem-related mechanisms operating outside conscious awareness, voluntary breathing is predicated on the individual’s conscious awareness and control of respiration and is a core component of yoga and other meditative practices (Anālayo, 2019; Peng et al., 2004). Some forms of voluntary respiration, such as slow breathing, are widely considered an important means of quickly reducing anxiety or stress (Balban et al., 2023; Birdee et al., 2023), whereas other forms such as rapid breathing (so called “breath of fire”) are sometimes considered to quickly increase energy levels (Peng et al., 2004). Despite increasing studies on the effects of voluntary breathing on the body and mind (Herrero et al., 2018; Zaccaro et al., 2018), the neural mechanisms underlying voluntary breathing control remain unclear.

Automatic breathing, controlled by the brainstem, maintains oxygen and carbon dioxide levels reflexively and independent of conscious control mechanisms. It is initiated by neurons in the formatio reticularis (FR), with the rhythm regulated by the dorsal respiratory group (DRG), the ventral respiratory group (VRG), and the pre-Bötzinger complex (preBötC) (Anderson & Ramirez, 2017; Shweta, Pramod & Chen, 2021; Ikeda et al., 2017; Yang & Feldman, 2018) (Figure 1b). The pneumotaxic and apneustic centers function to prevent lung over-distension. Automatic breathing can be interrupted or overridden by voluntary control mechanisms, and while there are separate anatomic pathways for automatic and voluntary breathing, these pathways demonstrate significant functional overlap (Prasad, Pal & Chen, 2021; Betka et al., 2022). Automatic breathing, controlled by FR’s respiratory networks, chemoreceptors, and respiratory mechanoreceptors, can be influenced by descending inputs from emotional and cognitive neural networks. Voluntary breathing thus appears to shift neural control towards supratentorial brain centers (Del Negro, Funk & Feldman, 2018; Homma & Masaoka, 2008; McKay et al., 2003; Herrero et al., 2018).

**Figure 1:**
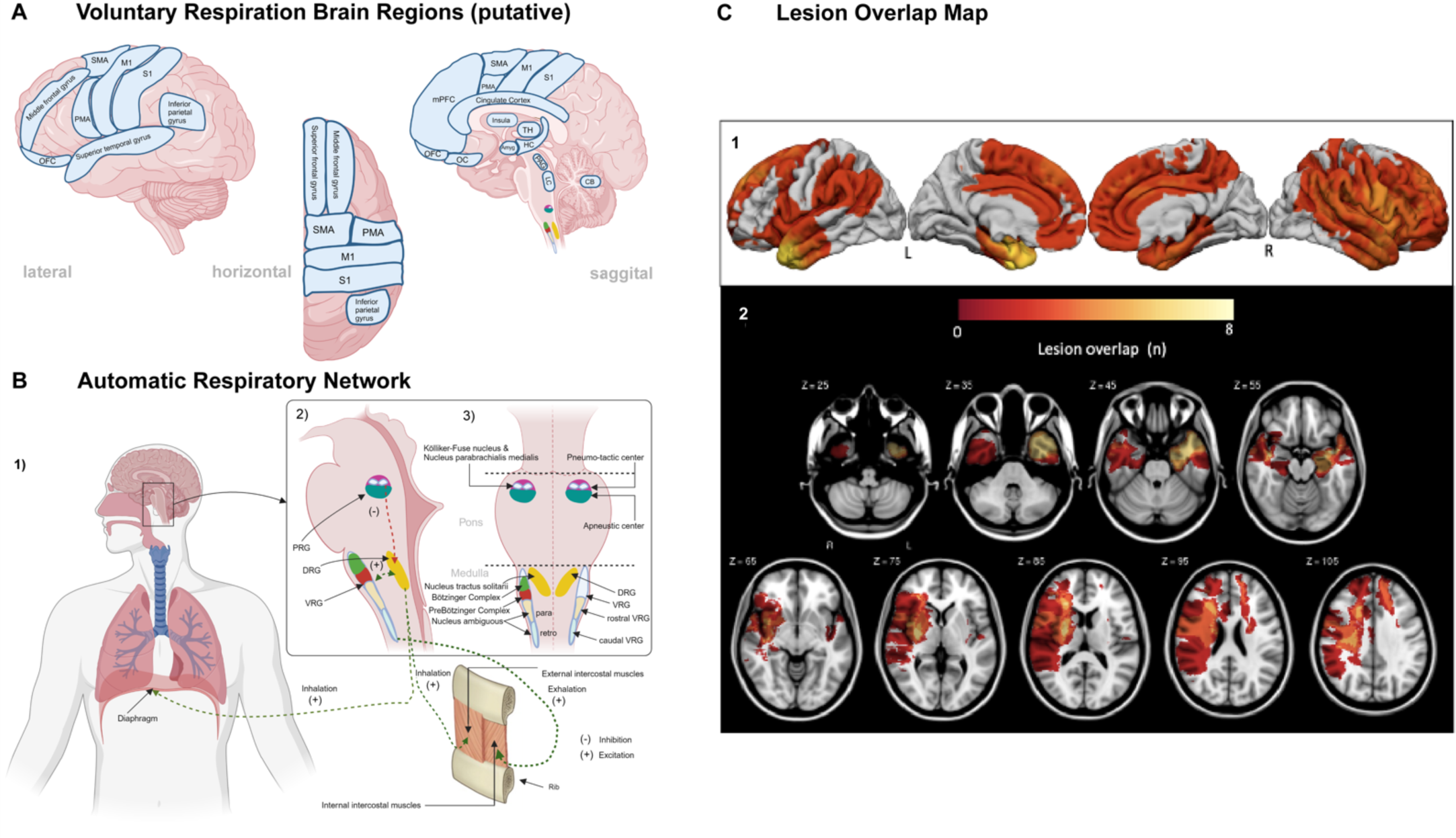
Overview of Brain Areas Involved in Voluntary and Automatic Respiration and Lesion Overlap Map for Participants with Brain Lesions. *Note.* **(A)** Location of human cortical areas putatively involved in the regulation of voluntary breathing. In addition to the respiratory centers of automatic breathing in the FR in the brainstem, several higher-order subcortical and cortical areas are likely involved in voluntary breathing. These are shown projected onto lateral, horizontal, and mid-sagittal surface views of the brain. The precise manner in which these regions control voluntary respiration is unknown. Abbreviations: Amyg – Amygdala, CB – Cerebellum, HC – Hippocampus, ILC - Infralimbic cortex, LC - Locus coeruleus, M1 - Primary motor cortex (1), OC - Olfactory cortex, OFC - Orbitofrontal cortex, PAG - Periaqueductal gray, PMA - Pre-motor cortex, SMA - Supplementary motor area, S1 - Sensory motor cortex (S1), TH – Thalamus. **(B)** Location of the major brainstem respiratory groups, projected onto the planes of a lateral sagittal section (2) and a horizontal section (3). The respiratory groups (right inset (2) and (3)) form a set of interacting structures: the Pontine Respiratory Group (PRG) in the superior part of the pons sends inhibitory signals to the Dorsal Respiratory Group (DRG) in the medulla and controls the filling phase of respiration. The DRG also receives signals from peripheral chemoreceptors and other receptors via vagus and glossopharyngeal nerves. These impulses produce inspiratory movements and are responsible for the basic rhythm of breathing. The Ventral Respiratory Group (VRG) causes either inhalation or exhalation. It is inactive during quiet breathing (i.e., it is widely accepted that exhalation is passive at rest), but it plays an important role in stimulating abdominal expiratory muscles during high respiratory demand. Abbreviations: DRG - Dorsal Respiratory Group, PRG - Pontine Respiratory Group, VRG - Ventral Respiratory Group. **(C)** This figure shows the cortical and subcortical overlap of lesions for the 20 lesion participants via lateral (1) and horizontal (2) views. Lesions were concentrated in the left temporal lobe, bilateral frontal lobe, the cingulate and insular cortices, and in subcortical structures such as the basal ganglia.

Functional neuroimaging and intracranial recording studies have identified frontal, parietal, and temporal lobe involvement in voluntary breathing during phasic increases in breathing rate (McKay et al., 2003; Herrero et al., 2018), suggesting a potential regulatory influence of widespread cortical and subcortical regions (Figure 1a). The prefrontal cortex (PFC), encompassing the dorsolateral, ventrolateral, and dorsomedial sections, connects to key subcortical respiratory regions (Jodo et al., 1998; Gabbott et al., 2005; Terreberry & Neafsey, 1987; Dutschmann & Dick, 2012), leaving it well positioned for a role in voluntary respiration in addition to its well-recognized roles in emotion regulation (Graham & Milad, 2013; Dixon et al, 2017; Silvers et al., 2015), motor control (Fine & Hayden, 2021; Watanabe et al., 2002), regulation of executive functions (Yuan & Raz, 2014; Friedman & Robbins, 2022), response inhibition (Aron, Robbins & Poldrack, 2014; Anderson, Bunce & Barbas, 2016), working memory (Curtis & Esposito, 2003; Lara & Wallis, 2015) and decision making (Bechara et al., 1998; Gläscher et al., 2012). The anterior cingulate cortex (ACC), another frontal region contributing to emotion (Etkin, Egner & Kalisch, 2011; Etkin, Büchel & Gross, 2015; García-Cabezas & Barbas, 2017), motivation (Holroyd & Yeung, 2012; Rolls, 2023), visceromotor regulation (Burns & Wyss, 1985) and cognitive control (Botvinick et al., 2001; García et al., 2022), has also been implicated in voluntary control of breathing. For example, using intracranial recordings in humans, a recent study linked dorsal ACC (dACC) activity with voluntary breath holding and increased blood carbon dioxide levels (Holton et al., 2021). The insular cortex (IC), a key region for processing and integrating interoceptive signals (Craig, 2002; Cameron, 2009; Khalsa et al., 2009a; Hassanpour et al., 2018) and emotion (Damasio et al., 2000), is engaged by expiratory breath holding (Pattinson et al., 2009), and right IC damage can lead to reduced dyspnea perception (Schön et al., 2008) and sleep-disordered breathing (Harper et al., 2021). The subcortical basal ganglia (BG) are also found to play a role in voluntary respiratory regulation (McKay et al., 2008) in addition to roles in initiating desired skeletomotor movement and inhibiting competing movements (Young, Reddy & Sonne, 2021; Martinu et al., 2012), as are the primary motor/sensory, premotor and supplementary motor areas. Temporal lobe damage can impact breathing regulation (Homma & Masaoka, 2008; Nobis et al., 2019), and patients with temporal lobe epilepsy may experience oxygen desaturation and central apnea during seizure-induced interference with amygdala activity (Dlouhy et al., 2015; Nobis et al., 2019; Rhone et al., 2020). In sudden unexplained death from epilepsy (SUDEP), studies have pointed toward a deficient arousal sensor to increased levels of CO2, which commonly occur during states of apnea (Buchanan, 2019), and a recent study even identified a subregion within the amygdala responsible for suppressing breathing and air hunger following stimulation (Harmata et al., 2023). Studies showing a role for the amygdala in interoceptive anxiety and cardiorespiratory interoception (Feinstein et al., 2013, Khalsa et al., 2016) have led to the suggestion that a lack of awareness of amygdala-mediated apnea may be a key mechanism provoking interoceptive arousal, fear, and anxiety (Feinstein, Gould & Khalsa, 2022). In summary, the intracranial and functional neuroimaging evidence to date suggests that the regulation of voluntary respiration is associated with neural processes spanning the PFC, ACC, insula, BG, amygdala, and adjacent temporal structures. However, there is a paucity of studies evaluating the impact of damage to these regions on voluntary breathing or associated affective, interoceptive, and autonomic processes.

The current study investigated the association of damage to human frontal, temporal, or limbic (Heimer & Van Hoesen, 2005) brain regions with the ability to sense and voluntarily regulate breathing patterns using a multilevel repeated measures design incorporating interoceptive, affective, and autonomic signals. We compared individuals with focal acquired brain lesions to healthy individuals, aiming to identify associations between cortical and subcortical damage and voluntary respiratory regulation, as well as to measures of physiological (i.e., autonomic) and subjective (i.e., affective and interoceptive) experience. To rule out general motor skill impairment, we included motor control tasks. We selected a number of symmetric respiratory rates, reflecting an interest in evaluating respiratory performance across the full physiological range of rates used in relaxation-, yoga-, and exercise-related behaviors, and to examine the impact of both slow and fast breathing on affective, interoceptive, and autonomic indices. We also selected an irregular breathing task to evaluate autonomic dynamics via respiratory sinus arrhythmia-induced heart rate variability and its impact on affective and interoceptive states. We hypothesized that individuals with damage to the aforementioned regions would show a reduced ability to regulate their respiratory performance during voluntary breathing, as indicated by decreased ability to follow a visually presented breathing trace in comparison to age-, sex-, and body mass index-matched healthy individuals without brain injury. Performance was evaluated with the peak cross-correlation between the instructed trace and chest wall excursion, measured with a chest belt. We hypothesized that any difference of performance would be specific to breathing, i.e., there would be no significant group differences in motor regulation ability on a hand movement task, due to the lesion locations in the selected group. With respect to affective and interoceptive indices, we hypothesized that the lesion group would report more anxiety across both slow and rapid breathing conditions versus resting rates, reflecting a U-shaped pattern of anxiety and task difficulty indicative of low levels of stress or anxiety at moderate levels of the stimulus (instructed breathing at normal respiratory rates) and higher levels of stress or anxiety at both the low and high extremes of the instructed breathing stimulus. Since rapid breathing conditions have been physiologically linked to stress and panic response, we expected that the increased demand on voluntary control to keep up with the fast pace would preferentially elevate anxiety levels in the lesion group. Although slow breathing techniques are often recommended for reducing anxiety and stress, we hypothesized that increased anxiety would occur in the lesion group due to a difficulty in keeping up with the demand for voluntary breathing regulation. Finally, we calculated multilevel correlations to explore associations between respiratory regulation parameters and aspects of autonomic signalling (i.e., skin conductance responses), affective response (i.e., anxiety ratings), and metacognitive perception (i.e., task difficulty ratings).

## Methods

### Participants

Healthy participants were recruited through public advertisements at the University of Iowa. Lesion participants with acquired focal brain injury were recruited from the Iowa Neurological Patient Registry at the University of Iowa Hospitals and Clinics. Inclusion criteria for the lesion participants included age 18 to 85 years, presence of a focal brain lesion at least 3 months after onset, no history of learning disabilities, psychiatric disorders, substance abuse, personality disorders, developmental epilepsy, or other unrelated neurological conditions. The lesions among the participants with focal brain injury were caused by ischemic or haemorrhagic strokes, meningioma resection, or surgical intervention for epilepsy. Exclusion criteria for the lesion participants included cardiovascular disease, chronic obstructive pulmonary disease, active substance use disorder, peripheral/central neurological disorders other than the defined brain lesion. Healthy participants were required to have no brain lesions and to meet the remainder of the inclusion/exclusion criteria.

### Participant Demographics

A total of 76 participants were consented, consisting of 56 healthy individuals (mean age (*M*) = 43.2, standard deviation (*SD*) = 12.5 years, range from 20 to 63 years) and 20 participants with acquired focal brain injury (*M* = 53.8, *SD* = 7.05 years, range from 36 to 84 years). Due to substantial variations in age and sex ratio between groups, the sample of 20 lesion participants (14 female and 6 male) was individually matched to an equal-sized sample of healthy participants (*n* = 20; 14 female and 6 male participants) based on age, sex, and body mass index (BMI; Table 1). The mean age of lesion (*M* = 53.8; *SD* = 7.05 years) and healthy (*M* = 53.6; *SD* = 10.6 years) participants did not significantly differ (*t*(33) = −0.07, *p* = 0.94), nor did the mean BMI of lesion (*M* = 27.6; *SD* = 5.0) and healthy (*M* = 30.7; *SD* = 5.9) participants significantly differ (*t*(35) = 1.81, *p* = 0.079). Shapiro-Wilk testing showed no evidence of non-normality of matching variables within experimental groups (Age: Lesion, *W* = 0.97, *p* = 0.734; Healthy, *W* = 0.92, *p* = 0.098; BMI: Lesion, *W* = 0.91, *p* = 0.071; Healthy, *W* = 0.97, *p* = 0.733).

**Table 1.** Demographic Characteristics of the Lesion and Healthy Samples.

| Variables | Lesion (n = 20) | Healthy (n = 20) | $\chi^2$ | t-ratio (df) | p-value |
| --- | --- | --- | --- | --- | --- |
| Age (years) |  |  |  | -0.07 (33) | 0.94 |
| Mean ± SD | 53.8 ± 7.1 | 53.6 ± 10.6 | | | |
| Sex (%) |  |  |  |  |  |
| Female | 70 | 70 |  |  |  |
| Male | 30 | 30 |  |  |  |
| BMI (kg / m<sup>2</sup>) |  |  |  | 1.81 (35) | 0.08 |
| Mean ± SD | 27.6 ± 5.0 | 30.7 ± 5.9 | | | |
| Median (IQR) | 26.7 (23.5 - 29.6) | 32.9 (27.1 - 34.5) |  |  |  |
| Handedness (n) |  |  | 2.11 |  | 0.15 |
| Right | 18 | 20 |  |  |  |
| Left | 2 | 0 |  |  |  |
| Education |  |  |  |  |  |
| Mean ± SD | 14.1 ± 1.9 | 16.6 ± 2.1 | | -3.8 (38) | <0.001 |
| Race (number) |  |  | 1.02 |  | 0.31 |
| White / Caucasian | 20 | 19 |  |  |  |
| Asian | 0 | 1 |  |  |  |
| Black | 0 | 0 |  |  |  |
| Hispanic | 0 | 0 |  |  |  |
| Mixed | 0 | 0 |  |  |  |

*Note.* Data shown compared by *t*-test or $\chi^2$ - test. BMI: Body mass index; IQR: Interquartile range; kg: Kilogram; m<sup>2</sup>: Square meter; n: Sample size; SD: Standard deviation;

### Brain Lesion Characteristics

The lesion locations for the 20 study participants with focal brain injury included a clustering of regions in the left and right frontal lobe, right hemispheric lesions in the insular cortex and basal ganglia, and left hemispheric lesions in the temporal lobe (Figure 1c).

### Experimental Design

After obtaining written informed consent participants completed a screening interview to verify eligibility. Inclusion and exclusion criteria for the lesion group were examined using medical history data from the Iowa Neurological Patient Registry. When applicable, new demographic information was also gathered.

Prior to completing the regulation tasks participants with brain lesions were asked to perform a neuropsychological motor coordination task, called the grooved pegboard test (Trites, 1989). The task involves fitting small irregularly shaped rods into openings on a pegboard using only one hand, providing a measure of the motor skills of the fingers, fingertips, hands, and forearms. This test of motor skill helped to identify fine hand-motor dysfunctions associated with acquired brain lesions, which could potentially confound the measurement of dial rotation in the hand motor regulation task. Since this measure was collected only in the lesion participants, their performance was compared against an age-matched normative sample (Lafayette Instrument Company, 2002). We employed individual scoring criteria, using Trites’ (1989) age-based norms and considered scores greater than 2 standard deviations (SD) above the age-based mean to reflect motor impairment (performance was measured via completion time, with higher durations indicating worse performance).

The main experimental condition included two regulation paradigms: respiratory and hand motor, each with two distinct blocks of tests (regular and irregular). For each paradigm, participants were seated and connected to measurement devices (see Measurements and Recording Devices section below). Each regular respiratory condition consisted of four 2-minute symmetric breathing trials at different instructed frequencies (6, 10, 16, and 60 nasal breaths per minute, bpm) and one irregular trial consisting of a pseudo-random frequency and amplitude modulation of the nasal inspiration and expiration phases. During the respiratory regulation trials, participants followed a visual cue with their breath, inhaling nasally with their mouth closed when a red dot moved up and exhaling nasally when it moved down a sinusoidal curve (Figure 2a). Prior to the respiratory task participants underwent an amplitude calibration test where they were instructed to demonstrate a maximum amplitude (i.e., full vital capacity) inhalation followed by exhalation. They were then told that the top and bottom of the screen represented their full vital capacity inhalation/exhalation, shown that the amplitude of the instructed sinusoid did not reach this boundary, and were told to ensure that the amplitudes of their inhalation/exhalations should be not exceed their comfort level. The hand motor task, which served as a control, involved the same visual cue, with participants instead making rotational dial movements with their dominant hand (or unaffected hemisphere/hand pair) to track the movement of the dot along the line. It included five trials with hand dial rotational movement cues that were matched to the respiratory task. Participants were also told not to twist their hands during the dial task to the point of discomfort. The presentation order of the experimental trials within each task was randomized, and the sequence of regulation tasks (breathing or motor first) was counterbalanced. Practice rounds were performed at the start of each block to familiarize participants with the experimental procedures.

**Figure 2:**
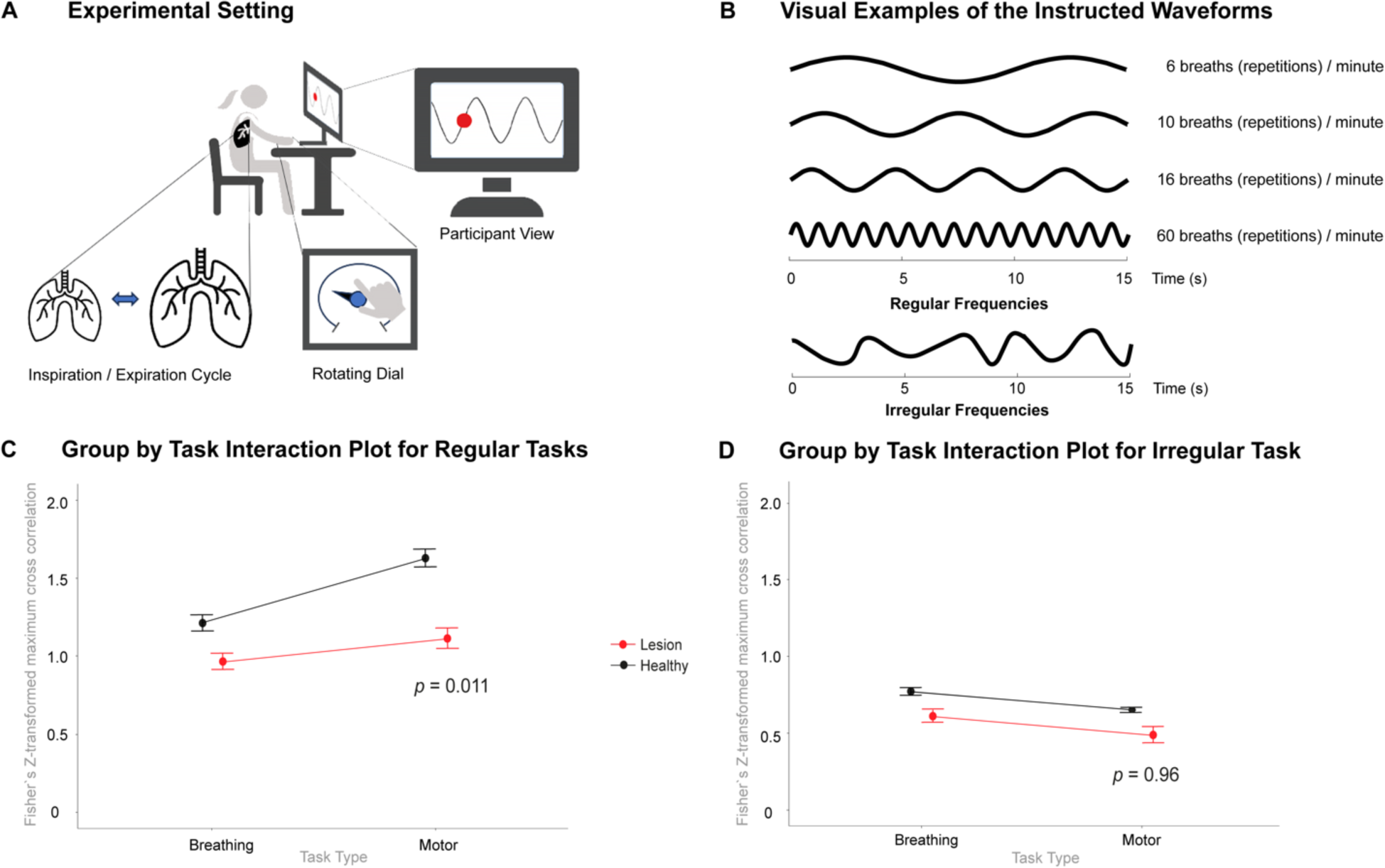
Overview of the Experimental Setup and Overall Performance for the Regular and Irregular Breathing and Motor Control Tasks. *Note.* **(A)** Participants were seated upright in front of a screen showing the corresponding instruction signal as a wave movement with a red dot moving up and down. **(B)** Exemplary 15-second illustration of instructed waveforms for regular and irregular experimental task conditions. For the regular version of the respiratory and motor regulation tasks, there were four experimental conditions ranging from 6 to 60 breaths (or repetitions) per minute. For the irregular task, both the amplitude and rate were randomly varied, with a frequency range from 6 to 60 breaths (or repetitions) per minute. Each experimental condition in the study lasted 2 minutes. Mean group performance across all conditions of the **(C)** regular task and **(D)** irregular task; performance was indexed via Fisher’s Z-transformed maximum cross correlation values (range: –2.65 to 2.65). P-values indicate the group by task interaction. Error bars indicate standard error.

After each trial, participants were asked to rate the physical sensations and emotional states they had experienced via a questionnaire involving multiple questions using a Likert response scale ranging from 1 (very little) to 5 (very much). This also included their perception of the difficulty and enjoyment of the task. Post-trial ratings of task difficulty were also evaluated to ascertain any differences in perceived difficulty of the respiratory and motor regulation tasks between the groups. They were also asked to rate other feelings and sensations experienced during each trial, including anxiety, boredom, focus, calm, excitement, and others. This study focused on evaluating task-related affect, specifically, anxiety and difficulty ratings.

### Measurements and Recording Devices

Demographic data (age, gender, handedness, ethnicity, and height) were gathered through a questionnaire. Body weight was physically measured to calculate BMI.

Physiological data were gathered via a MP100 acquisition unit (Biopac Inc., Santa Barbara, CA). Breathing activity was monitored using an elastic thoracic breathing belt and amplifier (RSP100C) at the mid-sternal level. Cardiac activity was measured using two electrodes paired with an amplifier (ECG100C) designed for electrocardiogram (ECG) monitoring and recorded at 200 Hz with a high pass filter to stabilize the ECG baseline. The positive electrode was positioned on the left lower anterior abdomen and the negative electrode on the right anterior chest at the mid-clavicular line (lead II configuration). Galvanic skin response electrodes, placed on the thenar and hypothenar eminences of the hand contralateral to the dial hand, were collected using a single-channel amplifier (EDA100C) at 200 Hz.

A custom-built dial was employed to measure hand movement regulation, consisting of a rotating potentiometer with a continuous rating scale ranging from 0.000 to 5.000 Volts at a sampling rate of 200 Hz, as utilized in our previous studies (Khalsa et al., 2009b; 2020). Participants were asked to rotate the dial using their dominant hand following a visual cue, while breathing normally during each trial.

### Statistical Analysis

Descriptive statistics for demographic data were compared for the lesion and matched-healthy comparison group. T-tests were used for metric variables (age, BMI), with normality checked using the Shapiro-Wilk test and histogram distributions.

To evaluate respiratory and motor regulation performance, maximum cross-correlation values were calculated between the instructed and measured signals (respiratory trace for breathing trials and dial trace for motor trials) using MatLab R2021a software (Mathworks Inc., Natick, MA) and transformed using Fisher’s Z-transformation to address skewed sampling distributions of correlation coefficients (Silver & Dunlap, 1987). Cross-correlations were calculated using the built-in xcorr function. Linear mixed effects models (LME) were used to compare performance across the respiration and motor regulation tasks via separate models for the regular and irregular tasks using the lme4 package in R (Bates et al., 2015).

For LMEs, random intercepts were included for each subject, as represented by the term (1|ID) in the model formulations; see next paragraph. This allowed us to include individual baseline values for each subject to account for between-participant variability. Additionally, we controlled for age in our models to account for its potential influence on the outcomes. However, random slopes for individual subjects were not included in these models, as it was assumed that the variability in slopes across subjects was not essential for the dependent variables under investigation.

We used the respective interaction terms between group membership (healthy vs. lesion) and the other predictor variables (breathing and motor performance, anxiety, HRV) as fixed effects for all tests, while controlling for age. The full term is outlined here as a model: value ∼ group*condition + age + (1|ID).

For the significance tests of the fixed effects in our Linear Mixed-Effects (LME) models, Wald chi-square tests were conducted. These tests aimed to determine the significance of the fixed effects and were carried out using the summary-function in conjunction with the *lmerTest* package in R (Kuznetsova, Brockhoff & Christensen, 2022). For the post-hoc analysis, the *emmeans* R-package (Lenth, 2018) was employed. The analysis excluded participants who were missing respiratory or motor regulation data for more than two trials per task. In the multilevel correlation analysis, Spearman’s correlation coefficients were calculated using R.

For skin conductance response (SCR) analysis, we utilized the open-source software Ledalab (version 3.x; Benedek & Kaernbach, 2010) for Matlab. Data were first preprocessed using a 1st order low-pass Butterworth filter with a cutoff frequency of 5 Hz. Adaptive smoothing was performed using Gaussian windows with a width of 16 samples. CDA.ISCR (Cumulative Driver Amplitude Integrated with Skin Conductance Response) was the dependent variable calculated for in-depth analysis of the SCR signals. CDA.ISCR represents the area covered by the temporal integration of the phasic driver of the SCR with respect to the duration of the SCR. This is achieved by multiplying the SCR by the size of the response window. CDA.ISCR is expressed in microsiemens-seconds (μS*s) and accounts for both the intensity and duration of the SCR over time. This allows a detailed study of the temporal evolution and intensity of responses to different stimuli.

### Ethics and Data Source

This study was approved by the University of Iowa Institutional Review Board. All volunteers provided written informed consent prior to study participation and received compensation for their involvement.

## Results

### Grooved Pegboard Task

For the nondominant hand, the lesion group mean completion time was *M* = 87.9 seconds (*SD* = 27.5); for the dominant hand, the mean score was *M* = 82.8 seconds (*SD* = 29.8). According to the scoring criteria, 29.4% (5 out of 17 participants) were classified as impaired for the non-dominant hand; for the dominant hand (i.e., the hand used to rotate the dial), 11.1% (2 of 18 participants) were classified as impaired.

### Respiratory and Motor Regulation Task Performance

Due to the putative widespread involvement of frontal, parietal, and temporal lobe structures in voluntary breathing, we conducted omnibus tests comparing performance of the entire lesion sample against the healthy sample for both the respiratory and motor regulation tasks. Due to the substantial dissimilarity of the irregular trial stimuli, we divided the evaluation of performance across the regular and irregular tasks.

### Regular Task

On the regular task there were significant main effects of group (*F*(1, 38) = 30.01, *p* < .001), task (*F*(1, 271) = 33.99, *p* < 0.001), and age (*F*(1, 37) = 6.77, *p* = 0.013) as well as a significant interaction of group and task (*F*(1, 271) = 6.19, *p* = 0.013). The age effect suggested that performance was worse with increasing age on both tasks. Post-hoc tests showed that the healthy comparison group significantly outperformed the lesion group across both breathing (*p* = 0.029, mean difference (*Mdiff*) = 0.24, 95%-CI [0.02, 0.47]) and motor control tasks (*p* < 0.001, *Mdiff* = 0.50, 95%-CI [0.26, 0.74]). The healthy comparison group performed significantly better on the hand motor control task as compared to the breathing task (healthy, *p* < 0.001, *Mdiff* = −0.44, 95%-CI [−0.63, −0.24]. See Figure 2c and Supplemental Figure 1 for performance at the individual task frequencies. In summary, while the lesion group performed significantly worse than the healthy comparison group on both regular breathing and regular motor tasks, the disparity in performance between the tasks was smaller in the lesion group as evidenced by the significant group by task interaction.

### Irregular Task

On the irregular task there was a significant main effect of group (*F*(1,37) = 14.16, *p* < 0.001) and task (*F*(1,37) = 17.00, *p* < 0.001). However, no significant main effect for age (*F*(1,37) = 0.16, *p =* 0.690) and no interaction was observed between group and task (*F*(1,37) = 0.002, *p* = 0.965). Post-hoc comparisons revealed that the healthy participants significantly outperformed those with lesions for both the breathing (*p* = 0.016, *Mdiff* = 0.16, 95%-CI [0.02, 0.3]) and motor tasks (*p* = 0.019, *Mdiff* = 0.16, 95%-CI [0.02, 0.3]). However, contrary to the pattern observed on the regular task, for the irregular task both groups performed significantly better on the breathing component of the task as opposed to the hand motor control (healthy, *p* = 0.037, *Mdiff* = 0.12, 95%-CI [0.00, 0.23]; lesion, *p* = 0.036, *Mdiff* = 0.12, 95%-CI [0.00, 0.24]). See Figure 2d and Supplemental Figure 1. In summary, participants with lesions demonstrated lower performance on both irregular tasks compared to healthy participants, with performance in the breathing condition being superior for both groups.

### Affective Ratings

For anxiety experienced during the regular breathing condition significant group differences were observed (*F*(1, 36) = 7.73, *p* = 0.009), with the lesion participants reporting higher anxiety (*M* = 2.02 out of 5, *SD* = 1.39) than the healthy comparisons (*M* = 1.39 out of 5, *SD* = 0.70) across all breathing tasks. There was also a significant effect of breathing condition (*F*(4, 147) = 14.48, *p* < 0.001), and an interaction between group and breathing condition (*F*(4, 147) = 3.16, *p* = 0.016). There was no main effect for age (*F*(1, 36) = 0.59, *p* = 0.446). Post-hoc tests revealed significant within-group differences in anxiety only at the 60 bpm condition relative to baseline for participants with lesions (*p* < 0.001, *Mdiff* = −1.91, 95%-CI[−2.77, −1.05]). This was also reflected by a significant difference in anxiety between the lesion versus healthy comparison group at 60 bpm (*p* = 0.001, *Mdiff* = 1.42, 95%-CI [0.32, 2.52]) (Figure 3a).

**Figure 3:**
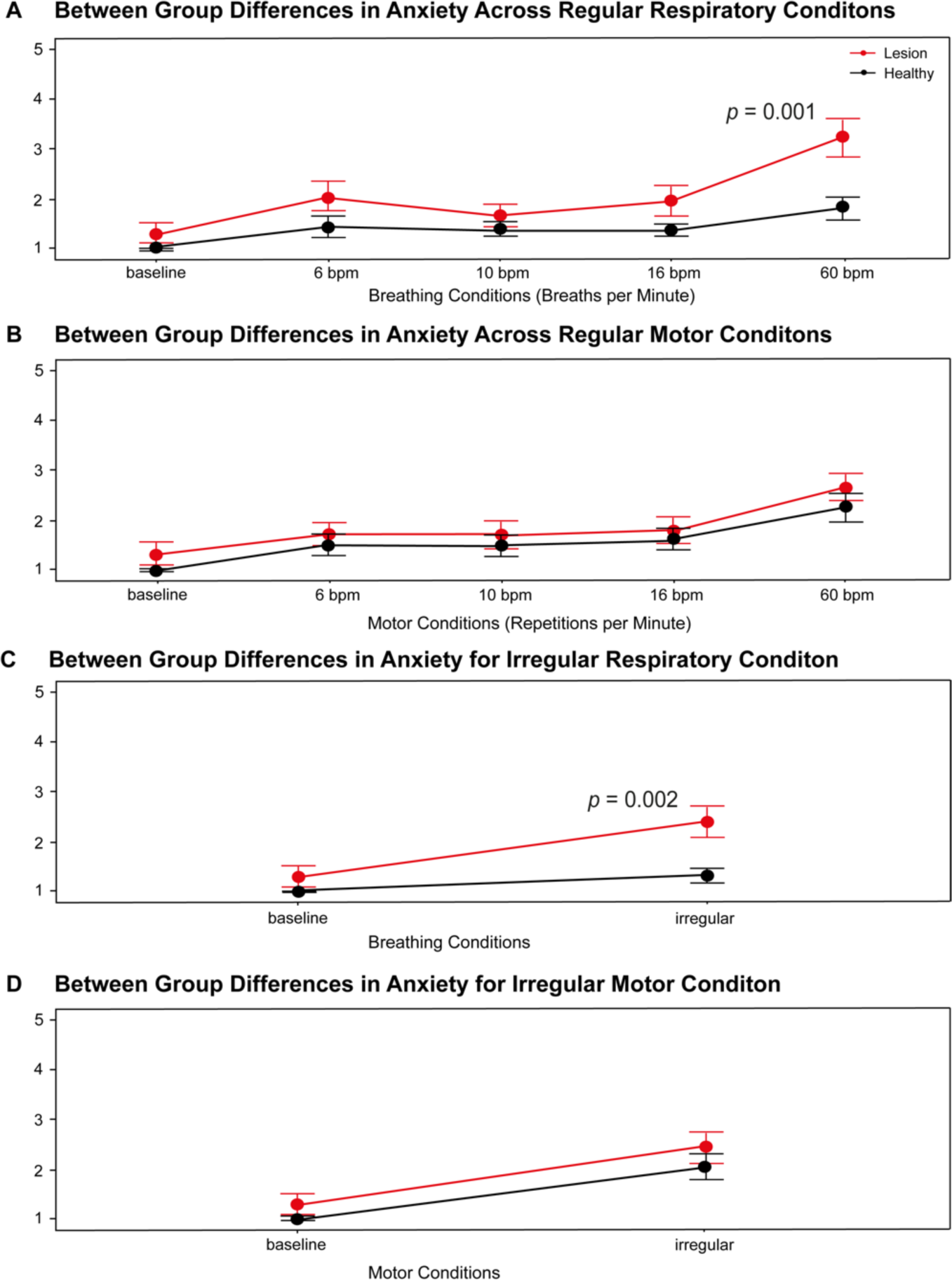
Mean Anxiety Ratings for All Task Types and Conditions *Note.* P-values indicate post-hoc between-group differences in anxiety for the 60-bpm condition of the regular breathing task and for the irregular breathing condition, calculated following observation of a significant group by condition interaction. Error bars indicate standard error.

For anxiety ratings during the regular motor task conditions, there was a significant main effect for task condition (*F*(4, 134) = 12.45, *p* < 0.001) but no significant main effect for the experimental groups (*F*(1, 34) = 1.34, *p* = 0.255) and age (*F*(1, 34) = 3.37, *p* = 0.075). Post-hoc tests showed that both groups reported significant anxiety increases for the fastest motor regulation trial (60 rpm) compared to baseline. Lesion participants also reported significant anxiety increases in slow and normal regulation trials (6 & 10 rpm), in the absence of significant group differences (Figure 3b).

For anxiety experienced during the irregular breathing condition, there was an interaction between group and breathing condition (*F*(1, 37) = 6.62, *p* = 0.014). Post-hoc tests showed significantly increased anxiety scores for the lesion versus healthy comparison group (*p* = 0.002, *Mdiff* = 1.08, 95%-CI [0.32, 1.85]) (Figure 3c).

For anxiety experienced during the irregular motor task, there was a significant main effect of task (*F*(1, 38) = 31.20, *p* < 0.001) but no significant main effect for the experimental groups (*F*(1, 38) = 1.97, *p* = 0.168), age (*F*(1, 38) = 0.09, *p* = 0.766) or interactions (Figure 3d).

### Difficulty Ratings

In assessing perceived difficulty during regular respiratory and motor tasks, there was a significant main effect for the groups (*F*(1, 36) = 9.67, *p* = 0.004) such that participants with lesions perceived the regular tasks as more challenging overall (Supplementary Figure 2). Additionally, there was a main effect for age (*F*(1, 35) = 7.04, *p* = 0.012), indicating that increased age was associated with perceived task difficulty. No significant interaction was found between group and task type (*F*(1, 262) = 0.10, *p* = 0.753). Post-hoc tests revealed that the healthy group had significantly lower difficulty ratings than the lesion group for the breathing task (*p* = 0.031, *Mdiff* = −0.74, 95% CI [−1.44, −0.045]), but not for the motor task (*p* = 0.090, *Mdiff* = −0.66, 95% CI [−1.38, 0.06]).

For the regular breathing conditions, there were no significant differences in the perceived difficulty between the breathing frequencies in the healthy subjects. However, the breathing rate of 60 bpm tended to be more difficult compared to 16 bpm (*p* = 0.124) and 6 bpm (*p* = 0.072). In the lesion group, the breathing frequency of 60 bpm was significantly more difficult compared to 10 bpm (*p* = 0.006) and 16 bpm (*p* < 0.001), while the differences between the other frequencies were not significant.

For the regular motor conditions, the healthy comparisons showed a significant increase in perceived difficulty at 60 rpm compared to 10 rpm (*p* < 0.001), 16 rpm (*p* = 0.006) and 6 rpm (*p* = 0.016). Similar results were observed in the lesion group, where 60 rpm was rated significantly more difficult compared to 10 rpm (*p* = 0.022), 16 rpm (*p* = 0.004) and 6 rpm (*p* = 0.026).

For the irregular tasks, there were no statistically significant differences in perceived difficulty between groups, either within one group (between breathing and motor conditions) or between groups across both tasks.

### SCR

For SCR during the regular breathing tasks, there was a main effect for the task (*F*(4, 136) = 5.70, *p* < 0.001). Post hoc analyses clarified that healthy participants in the 60 bpm showed significantly higher physiological arousal, than in the 6 bpm (*p* < 0.001, *Mdiff* = −6042.5, 95% CI[−10371, −1714]) and 10 bpm condition (*p* = 0.024, *Mdiff* = −4586.2, 95% CI[−8892, −280]). For lesion participants, there were no significant differences in SCR between the different respiratory tasks (see Supplementary Table 2).

For SCR during the regular motor tasks, there was also a main effect for the task (*F*(1, 122) = 2.97, *p* = 0.022). Post hoc analyses clarified that healthy participants in the 60 bpm showed significantly higher physiological arousal, than in the 6 bpm (*p* = 0.034, *Mdiff* = −5502, 95% CI[−10824, −181]). For lesion participants, there were no significant differences in SCR between the different motor tasks.

For the irregular breathing and motor tasks, there were no significant SCR effects for both experimental groups compared with baseline.

### Multilevel Correlation Analysis

This exploratory analysis examined the Spearman correlations between respiratory regulatory performance (behavioral/physiological measure), SCR (physiological measure), anxiety rating (subjective-affective measure), and rating of the respective perceived task difficulty (a metacognitive interoceptive measure). These were corrected for multiple comparisons via the Benjamini-Hochberg procedure. Based on our interest in the neural regulation of respiration, associations were evaluated for each group separately.

#### Correlations with Breathing

For healthy participants, significant correlations were observed across different conditions of the regular breathing task: At 10 bpm, a positive correlation was found between anxiety and perceived difficulty (*r*(20) = 0.70, *p* = 0.002) as well as between anxiety and SCR (*r*(20) = 0.65, *p* = 0.008) (Supplementary Table 2). At 16 bpm, anxiety was negatively correlated with performance accuracy (*r*(20) = −0.52, *p* = 0.030), and positively correlated with perceived difficulty (*r*(20) = 0.66, *p* = 0.048). In addition, a significant positive correlation was also found between perceived difficulty and SCR (*r*(20) = 0.58, *p* =0.029). There were no significant correlations found between variables in the slow breathing (6 bpm), fast breathing (60 bpm) or the irregular breathing task.

For lesion participants, significant correlations were observed across different conditions of the regular breathing task: At 16 bpm, anxiety was negatively correlated with performance accuracy (*r*(20) = −0.09, *p* = 0.016). Additionally, a significant positive correlation was found between perceived difficulty and skin conductance (*r*(20) = 0.42, *p* = 0.040). At 60 bpm, a significant negative correlation was found between performance accuracy and SCR (*r*(20) = −0.62, *p* = 0.039). Another significant positive correlation was found between anxiety and perceived difficulty (*r*(20) = 0.67, *p* = 0.020). There were no significant correlations observed for the irregular breathing task.

#### Correlations with Hand Movement

For healthy participants, significant correlations were observed across different experimental conditions of the motor dial task: At 6 rpm, anxiety was positively correlated with perceived difficulty (*r*(16) = 0.58, *p* = 0.009). No significant correlations were identified for the experimental trials at the 10, 16 and 60 rpm frequencies, nor were any found for the irregular motor task.

For lesion participants, the only significant correlation were observed at 60 rpm, between anxiety and perceived difficulty (*r*(16) = 0.77, *p* = 0.009). No significant correlations were identified for the experimental conditions for the experimental trials at the 6, 10 and 16 rpm frequencies, nor were any found for the irregular motor task.

### Exploratory Analyses

Given previous studies demonstrating breathing frequency-specific changes in heart rate variability (HRV), we conducted an exploratory analysis of HRV across all tasks. This analysis revealed no group differences across the different instructed respiratory frequencies but found that the lesion group (relative to healthy comparisons) exhibited slightly higher heart rates as well as lower low-frequency and higher high-frequency power of the HRV during the hand movement task (see Supplementary Table 3 and Supplementary Figures 3-6).

To evaluate the potential association of specific cortical networks with voluntary breathing control, we performed a detailed functional network analysis using each lesion participant’s lesion as a seed within a normative connectome to create lesion-related connectivity maps (Joutsa et al., 2023). This lesion network analysis showed that lesions tended to be grouped in approximately four of the Yeo7 networks (Yeo et al., 2015), namely the limbic network, default mode network, somatomotor network, and ventral attention network (Supplementary Figures 7-9).

## Discussion

The current study investigated the association of damage to cortical and subcortical brain areas with human respiratory and motor regulation, characterizing the interoceptive and affective responses occurring during with these tasks. As hypothesized, participants with frontal, temporal, and limbic brain lesions exhibited significantly lower performance at regulating their breathing across regular and irregular tasks, in addition to reductions in hand motor control. While both groups performed better on motor control tasks compared to breathing tasks during regular instruction conditions, this discrepancy was less pronounced among the lesion group. Moreover, lesion participants reported heightened anxiety, especially during rapid or irregular breathing tasks, which was not seen during hand motor tasks. This suggests an anxiogenic response specific to breathing challenges in the group with brain lesions.

A fundamental, yet under-researched, aspect of human brain-body interactions is the connection between the voluntary control of breathing and cortical and subcortical regions of the central nervous system. Here, we examined an omnibus scenario by testing a group of lesion patients with damage to a heterogeneous set of cortical and subcortical areas and comparing them to healthy participants. Our hypothesis that participants with damage to the frontal, temporal or limbic regions would show a reduced ability to control their breathing performance during a voluntary breathing task was partially supported. Indeed, the participant group with lesions was found to have, on average, a lower respiratory regulatory capacity than the healthy participant group across both regular and irregular instruction conditions. However, against our expectations, this reduction in performance was not restricted solely to breathing, suggesting that damage to the observed regions was also associated with other forms of rhythmic motor control. This inference is supported by the fact that the lesion group did not show substantial impairment in other forms of non-rhythmic hand movement, as measured by the grooved pegboard test, in relation to a normative age-matched reference sample (as we did not collect pegboard test data in the healthy comparator sample, we are unable to test directly for group differences on the task and cannot establish whether group differences would have been observed, which may be considered a limitation of the study). One implication for this finding is that the neural circuits for production of complex skeletomotor movements are recruited, and may even be necessary, for the optimization of voluntary breathing control patterns, such as deep breathing and rapid breathing, that are practiced during yoga, meditation, and stress management routines. While there is evidence from structural and functional brain imaging approaches supporting the engagement of temporal and frontal brain regions by yoga (Gothe et al., 2019), this is a sparsely studied topic and more data are needed.

Although both groups performed better on motor control tasks as compared to breathing tasks during regular instruction conditions, this discrepancy was less pronounced among the lesion group. The exact mechanisms underlying this phenomenon warrant further investigation, but the finding may suggest a specialized impairment of respiratory (versus motor) control in association with injury to the involved frontal, limbic, and temporal brain regions. At the same time, the non-significant interaction between group and task in the irregular instruction condition suggests a lower association with breathing control mechanisms when respiratory frequency and amplitude is randomly fluctuating. Respiratory rhythms vary considerably depending on the internal and external environmental context, with symmetric changes more common during homeostatic perturbations in the sympathetic (e.g., hyperventilation during fight or flight response) and parasympathetic (e.g., slow breathing during relaxation) domains. Accordingly, it seems plausible that the lesion group’s difficulty in symmetric respiratory regulation could manifest via tachypnea-associated anxiety during the 60-bpm condition; a possibility that is further supported by their greater respiratory-specific difficulty ratings.

The lack of respiratory/hand-motor discrepancy for irregular breathing conditions raises several considerations worth exploring. First, respiratory rhythms are increasingly acknowledged to influence the balance of excitatory and inhibitory activity within the brain, with a particular influence of shifts in excitability that are inducible by respiratory phase (Kluger & Gross, 2021; Kluger et al., 2023). This suggests that aperiodic changes in respiratory rhythm, such as those produced during the irregular breathing condition, could alter the brain’s excitatory/inhibitory coupling in a manner different from the regular breathing condition. Why these “respiration-modulated brain oscillations” (RMBOs) would lead to a relative reduction in breathing performance in the regular vs. irregular breathing condition is uncertain, but future studies might evaluate the impact of excitatory vs. inhibitory input from cortical and subcortical areas to respiratory brainstem nuclei. Such an approach would require an animal model of respiratory control, although neurosurgical patients could be another means of gaining some insight into the role of specific subregions (e.g., medial temporal regions in epilepsy patients awaiting a corrective temporal resection procedure). Second, respiratory rhythms are well established to influence cognition, affect, and perception through influences on olfactory, insular, and somatosensory pathways (reviewed in Allen, Varga & Heck, 2023) that are closely anatomically connected to the frontal, limbic, and temporal regions damaged in the present lesion sample. It seems plausible that the volatility inherent to the irregular respiration condition could have altered the excitability of cortical circuits in a manner different from the regular respiration conditions, but the precise impact of this volatility (e.g., selective influence on interoceptive prediction error processing, or global influence on exteroceptive or affective representations; Brændholt et al., 2023) is unknown and only the subject of speculation. Third, the present task relied primarily on the voluntary top-down control of motor circuitry. Adapting this approach to incorporate spontaneous breathing perturbations (e.g., via transient inspiratory restrictions (Herzog et al., 2021; Burt et al., 2023)) would allow for the discernment of bottom-up processes contributing to the interoceptive and affective differences reported during specific forms of breathing. Adapting such a protocol to the concurrent measurement of neural oscillations would facilitate the identification interactions between top-down and bottom-up processing streams, which could be used to develop fine-grained explanatory models of respiratory sensation and regulation. Incorporating a network perspective could be important to such approaches, as our exploratory functional network analysis of breathing performance suggested that lesions affecting broader limbic, default mode, somatomotor, and ventral attention networks could play a role in breathing control. Additional analytic approaches could also statistically evaluate correlations between lesion-related networks and behavior (Joutsa, Corp & Fox, 2022; Pustina et al., 2018), though such approaches typically require a larger sample size than that employed in the current study.

The affective responses observed during different task conditions provide intriguing insights into the interplay between brain lesions, task performance, and affective responses. First, the fact that the lesion group reported greater anxiety levels overall during the breathing task, but not during the motor task or at baseline, suggests a degree of aversive sensitivity to respiratory signals in general. Second, the fact that the lesion participants also reported higher anxiety levels during specific respiratory conditions, notably during fast breathing, highlights a potential heightened respiratory-affective sensitivity or alternatively a decreased capacity to emotionally regulate tachynpeic states following brain injury. Although there is no pre-lesion data from which to make firm claims of causality, the fact that the anxiety increase was specific to one breathing frequency band increases the likelihood that reduced neural resources post-injury might render these individuals more susceptible to interpreting tachypnea as an anxiety-inducing stimulus. This group also reported anxiety during the irregular breathing task, which spanned both rate- and amplitude-modulations, raising the possibility of additional factors such as rapid variability or unpredictability of breathing signals that could influence affective experience. Finally, the lack of anxious responses to slow breathing might be interpreted to suggest that individuals with brain lesions in frontal, limbic, and temporal areas may be able to utilize slow breathing techniques as taught for reducing anxiety and stress, without concern for a paradoxical effect.

In general, the correlations between regular task performance, anxiety, perceived difficulty, and physiological measures (such as SCR) showed similar patterns for both groups, although no formal statistical tests were conducted to compare the correlations between groups. For example, in both groups breathing performance accuracy was associated with lower difficulty ratings, greater difficulty was associated with greater SCRs, and greater difficulty was associated with greater anxiety. During the hand motor task there were fewer significant correlations for both groups. Collectively, these data suggest that increased anxiety might be detrimentally associated with breathing regulation ability irrespective of brain injury status. However, it is also possible that reduced breathing regulation ability is associated with anxiety at greater task difficulties. Future studies could examine whether these associations are altered in clinically anxious patients. The correlations between affective states and task performance observed during the hand movement task also illustrate the challenging nature of the task instruction. In healthy participants, anxiety was strongly correlated with perceived difficulty at the slowest rotation (6 rpm), while in lesioned participants, a negative correlation between the two was observed at the highest speed (60 rpm). Such rate-related variations could imply different coping or regulatory mechanisms were activated depending on the neural integrity and the nature of the task. The lack of significant correlational patterns for either group on the irregular breathing condition is unclear.

This study has several limitations and future directions. These include a small sample size, which was limited by the rarity of the lesion sample, and the heterogeneity of lesion location, which was determined by random sampling from available participants visiting the laboratory during the data collection period. Future studies should recruit larger and more homogeneous samples and would benefit from focal targeting of overlapping lesion locations to rigorously test hypotheses regarding neurorespiratory coupling. As study measurements were taken from patients during the chronic injury phase, it is possible that there were undetected impacts of injury on respiration or motor ability during the acute injury phase that had been compensated for by neural reorganization by the time of the study. Respiration was only measured using a thoracic chest belt, which can yield inaccurate measurements, thus future studies would benefit from more sophisticated measurement approaches capable of assessing variables such as tidal/vital capacity, nasal flow rates, and capnometry to index exhaled gas concentrations. Additionally, this task examined a restricted form of respiratory control; specifically, regular or irregular breathing across a range of respiratory frequencies (and in the latter task also across various respiratory depths). As we did not examine other forms other forms of breath control, such as breath holding (inspiratory or expiratory), valsalva maneuver, or respiratory pauses (such as those that may occur during speaking or singing), we cannot extend our inferences to these respiratory behaviors. The groups also differed with respect to educational level; although there are no clear suggestions this difference may have influenced the current results future studies might consider controlling for this variable. Future research directions include mechanistically investigating the influence of activity within specific brain regions on respiratory and motor regulation with larger, more homogeneous sample groups; including more cases of bilateral lesions; and possibly employing animal models and human intracranial recording studies to further understand the neural basis of respiratory and motor control.

### Conclusion

This study demonstrates that damage to frontal, temporal, or limbic brain regions is associated with abnormal voluntary respiratory and motor regulation and tachypnea-related anxiety, highlighting the role of the forebrain in affective and motor responses during breathing.

## Author Contributions

SSK, CK, and DT: conceived and designed the experiments; SSK, CK, HB: carried out the experiments and/or processed the data. SSK, SK, CK, JB, DT and HB: contributed to the analysis of the results. JB and HB: contributed the illustrations. Figure 1A and B was created with Biorender by HB and Figure 1C was created by JB, with input from SSK. All other figures were created by HB with input from SSK. SSK and HB: wrote the paper, with input from all authors.

