## Supplement for "Sensing, Feeling, and Regulating: Investigating the Association of Focal Brain Damage with Voluntary Respiratory and Motor Control"

**Table of contents**

**Supplementary Methods. Heart Rate Variability (HRV) Estimation…….…………………...............Page 2**

**Supplementary Results. Baseline and Task-Related Physiological Responses ………………………Page 2**

**Supplementary Results. HRV Analysis……………………………….…………………………………................Page 2**

**Supplementary Results. Functional Network Analysis…...…….………………………………….............Page 5**

**Supplementary Table 1. Baseline and Task-Related Physiological Responses** ***……………..*.........Page 6**

**Supplementary Table 2. Multilevel Task Correlation Matrices*.……………………*……………............Page 7**

**Supplementary Table 3. HRV Parameters in Time and Frequency Domains*…….*……................Page 8**

**Supplementary Figure 1. Respiratory/Hand Motor Regular Task Performance…....................Page 9**

**Supplementary Figure 2. Task Difficulty Ratings…………………………………………………………………..Page 10**

**Supplementary Figure 3. HRV Time Domain Parameter Distributions (Regular Tasks)............Page 11**

**Supplementary Figure 4. HRV Time Domain Parameter Distributions (Irregular Task)............Page 12**

**Supplementary Figure 5. HRV Frequency Domain Parameter Distributions (Regular Tasks)….Page 13**

**Supplementary Figure 6. HRV Frequency Domain Parameter Distributions (Irregular Task)….Page 14**

**Supplementary Figure 7. Spatial Correlation Patterns Between Brain Lesions and Yeo7 Network Domains….……………………………….……………………………………………….….……………………………………..Page 15**

**Supplementary Figure 8. Scree Plot of Principal Component Analysis of Cortical Networks….Page 16**

**Supplementary Figure 9. Spatial Visualization Principal Component Distribution…………………Page 17**

**Supplementary References………………………………….……………………………………………….….………….Page 18**

**Supplementary Methods. Heart Rate Variability (HRV) Estimation**

HRV parameters in the time- and frequency domain were calculated using MatLab *HRV Tool* (Vollmer, 2019; https://marcusvollmer.github.io/HRV/). Using the batch processing loop to analyze the group data, Average Heart Rate (AHR), RMSSD, pNN50 and SDDN were first calculated as HRV indicators for the time domain. For the frequency domain, HF- and LF-power were determined as HRV indicators using the fast Fourier transform and the available functions *HRV.fft_val* and *HRV.fft_val_fun*.

First, the recorded heart rate curves were visually examined for measurement artefacts such as repeated zero values and peak values > 200 bpm. No subject was excluded from further analysis. Raw data from each subject were plotted as RR interval-time graphs using the batch processing function (*batchprocessing.m*), i.e., the intervals of two consecutive heartbeats or R-spikes in the ECG were plotted in ms over the total time of 2 min per respiratory condition. For artifact reduction, the HRV.RRfilt.m function was used, which removes artifacts from RR sequences by relative RR intervals. Trend adjustment was not performed. For each 2-min interval, different HRV parameters were recorded in the categories of time domain and frequency domain by default according to the recommendations of the Task Force (1996). Due to 10% missing values for the AHR, RMSSD and SDNN in the motor regulation tasks, imputation of the missing data was modelled by multiple imputation using *mice* (Van Buuren & Groothuis-Oudshoorn, 2011).

To determine the frequency parameters, the RR interval-time data were converted to a continuous function in the HRV tool by spline interpolation. This was done using the nonparametric fast Fourier transform (FFT). Frequency components were defined according to the Task Force (1996) as follows: LF component 0.04-0.15 Hz and HF component 0.15-0.4 Hz.

For the statistical evaluation of the HRV parameters, group comparisons were made between the lesion participants and the healthy participants, after averaging across all trials for the regular task conditions. First, Levene's test was used to check the homogeneity of variances. Depending on the result of this test, the unpaired t-test (for equal variances) or Welch's test (for unequal variances) was applied to examine significant differences between the groups. The above tests were performed separately for the different HRV parameters, both in the time and frequency domains. All statistical analyses were performed with a significance threshold of p < 0.05.

**Supplementary Results. Baseline and Task-Related Physiological Responses**

The results of the Welch's Two Sample t-test for respiration over a period of two minutes showed no statistically significant differences between the groups, *t*(37) = -1.19, *p* = 0.243 (The situation was similar for the variable "SCR", where no significant differences were found, *t*(35) = 0.22, *p* = 0.826. There was also no significant difference between the groups for heart rate ("HR"), *t*(27) = -1.62, *p* = 0.116. In addition, the variables "SDNN" (*t*(18) = -0.62, *p* = 0.545), "RMSSD" (*t*(20) = -0.25, *p* = 0.804), "pNN50" (*t*(31) = 0.10, *p* = 0.924), "LF" (*t*(35) = 0.82, *p* = 0.416) and "HF" (*t*(35) = -0.82, *p* = 0.416) showed no statistically significant differences between the two groups. In summary, it can be said that breathing in healthy individuals and individuals with lesions shows no statistically significant differences with regard to the physiological variables measured during the baseline measurement.

**Supplementary Results. HRV Analysis**

We tested whether HRV (time and frequency domain) differed significantly between participants with lesions compared to healthy individuals during the regular and irregular respiratory regulation tasks and motor control tasks.

*Time Domain Analysis*

In the time domain of HRV analysis for the regular breathing tasks, we tested average heart rate (AHR), pNN50, RMSSD and SDNN for the respiratory regulation condition. A Welch's test revealed a statistically significant difference between the AHR values of the lesion group (*MD* = 75.43, *SD* = 14.26) and the healthy control group (*MD* = 71.36, *SD* = 8.59), with the AHR control group being 4.08 bpm lower on average (95%-CI[0.39, 7.76]), *t*(130) = 2.19, *p* = 0.003, *d* = 0.35. For RMSSD, subsequent testing by unpaired students t-test revealed no statistically significant difference between the RMSSD of the lesion group (*MD* = 24.28, *SD* = 15.31) and the healthy control group (*MD* = 25.19, *SD* = 12.85), *t*(158) = -0.41, *p* = 0.683. For pNN50, an unpaired students t-test revealed no statistically significant difference between the pNN50 of the lesion group (MD = 5.88, SD = 9.21) and the healthy control group (*MD* = 7.03, *SD* = 9.49), *t*(158) = -0.78, *p* = 0.438. Lastly, we tested SDNN parameters. An unpaired students t-test showed no statistically significant difference between the SDNN of the lesion group (*MD* = 33.11, *SD* = 17.43) and the healthy control group (*MD* = 37.05, *SD* = 18.71), *t*(158) = -1.37, *p* = 0.171.

For the regular motor regulation task, the results were as follows. For AHR, Welch's test revealed no statistically significant difference between the AHR scores of the lesion group (*MD* = 74.12, *SD* = 10.40) and the healthy control group (*MD* = 71.85, *SD* = 8.36), *t*(158) = 1.51, *p* = 0.134. Testing for RMSSD group differences with an unpaired Student's t-test revealed no statistically significant difference between the RMSSD of the lesion group (*MD* = 23.33, *SD* = 14.42) and the healthy control group (*MD* = 22.76, *SD* = 12.19), *t*(158) = 0.26, *p* = 0.793. Moreover, subsequent testing by unpaired students t-test revealed no statistically significant difference between the pNN50 of the lesion group (*MD* = 6.68, *SD* = 9.31) and the healthy control group (*MD* = 5.75, *SD* = 7.90), *t*(158) = 0.67, *p* = 0.503. Finally, we tested the SDNN parameters. Unpaired student's t-test revealed no statistically significant difference between the SDNN of the lesion group (*MD* = 27.57, *SD* = 11.27) and the healthy control group (*MD* =30.66, *SD* = 16.00), *t*(158) = -1.40, *p* = 0.163.

In the time domain of HRV analysis for the irregular breathing task, Welch's test revealed no statistically significant difference between the AHR values of the lesion group (*MD* = 73.67, *SD* = 13.23) and the healthy control group (*MD* = 71.17, *SD* = 7.81), *t*(37) = 0.47, *p* = 0.472. For RMSSD, subsequent testing by unpaired students t-test revealed no statistically significant difference between the RMSSD of the lesion group (*MD* = 26.34, *SD* = 13.82) and the healthy control group (*MD* = 30.78, *SD* = 13.67), *t*(38) = -1.02, *p* = 0.314. For pNN50, an unpaired students t-test revealed no statistically significant difference between the pNN50 of the lesion group (MD = 8.17, SD = 9.55) and the healthy control group (*MD* = 10.41, *SD* = 11.55), *t*(37) = -0.67, *p* = 0.509. Lastly, we tested SDNN parameters. An unpaired students t-test showed no statistically significant difference between the SDNN of the lesion group (*MD* = 31.19, *SD* = 12.74) and the healthy control group (*MD* = 37.03, *SD* = 13.56), *t*(38) = -1.40, *p* = 0.169.

For the irregular motor regulation task, the results were as follows. For AHR, Welch's test revealed no statistically significant difference between the AHR scores of the lesion group (*MD* = 73.58, *SD* = 10.08) and the healthy control group (*MD* = 71.78, *SD* = 8.74), *t*(34) = 0.58, *p* = 0.567. Testing for RMSSD group differences with an unpaired Student's t-test revealed no statistically significant difference between the RMSSD of the lesion group (*MD* = 39.80, *SD* = 31.48) and the healthy control group (*MD* = 34,73, *SD* = 42.63), *t*(35) = 0.41, *p* = 0.684. Moreover, subsequent testing by unpaired students t-test revealed no statistically significant difference between the pNN50 of the lesion group (*MD* = 7.89, *SD* = 11.78) and the healthy control group (*MD* = 5.67, *SD* = 11.52), *t*(35) = 0.57, *p* = 0.578. Finally, we tested the SDNN parameters. Unpaired student's t-test revealed no statistically significant difference between the SDNN of the lesion group (*MD* = 38.91, *SD* = 29.39) and the healthy control group (*MD* =37.09, *SD* = 30.41), *t*(35) = 0.19, *p* = 0.854.

*Frequency Domain Analysis*

In the frequency domain of HRV analysis for the regular breathing task, there were no significant group differences. Welch's test showed no significant difference between the LF of the lesion group (*MD* =54.28, *SD* = 27.09) and the healthy control group (*MD* =59.74, *SD* = 28.66), *t*(158) = - 1.28, *p* = 0.217, and for the HF between the lesion group (*MD* =45.72, *SD* = 27.09) and the healthy control group (*MD* =40.26, *SD* = 28.65), *t*(158) = -1.28, *p* = 0.217.

For the frequency parameters in the regular motor tasks, however, there were significant group differences. Welch's test showed significant difference between the LF of the lesion group (*MD* =47.99, *SD* = 23.05) and the healthy control group (*MD* = 55.58, *SD* = 19.80), *t*(154) = -2.28, *p* = 0.024, *d* = 0.36, and for HF between the lesion group (*MD* = 52.00, *SD* = 23.05) and the healthy control group (*MD* = 44.42, *SD* = 19.80), *t*(154) = 2.28, *p* = 0.024, *d* = 0.36. It was tested that for both the respiratory and motor regulation conditions, there were no significant differences between the experimental groups in terms of HRV parameters in the time domain and frequency domain, as can be seen from Supplementary Figure 3 and Figure 5. Only for the AHR difference between the groups in the respiratory task did we find a small effect and for the frequency parameters LF and HF in the motor task effects of medium size.

In the frequency domain of HRV analysis for the irregular breathing task, there were no significant group differences. Welch's test showed no significant difference between the LF of the lesion group (*MD* =51.11, *SD* = 20.49) and the healthy control group (*MD* = 57.23, *SD* = 16.81), *t*(37) = - 1.03, *p* = 0.309, and for the HF between the lesion group (*MD* =48.89, *SD* = 20.49) and the healthy control group (*MD* =42.77, *SD* = 16.81), *t*(38) = 1.03, *p* = 0.301.

For the frequency parameters in the irregular motor task, there were also no significant group differences. Welch's test showed no significant difference between the LF of the lesion group (*MD* =44.13, *SD* = 21.48) and the healthy control group (*MD* = 52.09, *SD* = 23.67), *t*(34) = -1.01, *p* = 0.319, and for HF between the lesion group (*MD* = 55.87, *SD* = 21.49) and the healthy control group (*MD* = 47.91, *SD* = 23.67), *t*(34) = 1.01, *p* = 0.319. It was tested that for both irregular, respiratory, and motor regulation condition, there were no significant differences between the experimental groups in terms of HRV parameters in the irregular time domain and frequency domain, as can be seen from Supplementary Figure 4 and Figure 6.

Breathing and autonomic function are closely linked, which makes the interpretation of the results complex. We do not have direct evidence that autonomic functions differed grossly between the groups, based on the lack of group differences in HR, RR, SCR, or HRV at baseline, although we did not assess for potential neural alterations that have been previously associated with autonomic function (Turchi et al., 2018; Barber et al, 2023 ). Breathing can also influence the autonomic signals that are reflected by HRV, to potentially attenuate or amplify HRV data. For example, changes in breathing associated with respiratory sinus arrhythmia (RSA), also can have a direct effect on the HRV signal. With controlled breathing, especially if it is rhythmic and slowed at 6 bpm, can be so strong that it results in distinct changes in certain spectral components of HRV (e.g., vagally-mediated low-frequency power, see Kromenacker et al., 2018 Psychosom Med ). In this respect, we cannot rule out the possibility that general respiratory and/or autonomic alterations producing reduced ascending arousal may also contributed to observed group differences in HRV during the motor task. We are less certain as to why the groups did not show differnces in HRV on the respiratory task.

**Supplementary Results. Functional Network Analysis**

To address the question of the importance of specific brain networks in voluntary respiratory control, we conducted a detailed functional network analysis, where each participant's lesions served as a starting point within a normative connectome to create lesion-related connectivity maps. These maps were then compared with the Yeo7 atlas to determine spatial correlations between the lesions and the networks defined in the atlas.

First, we created individual connectivity maps for each participant by using their brain lesions as starting points within a normative connectome, based on the GSP1000 dataset. This method allowed us to precisely map the effects of the lesions on the brain network. The created maps were then compared with the Yeo7 atlas, which divides the brain into seven functional networks. Through this comparison, we were able to determine which specific networks were affected by the lesions.

The analysis showed that lesions tended to be grouped in approximately four of the Yeo7 networks: Limbic Network (LN), Default Mode Network (DMN), Somatomotor Network (SMN), and Ventral Attention Network (VAN). This finding suggests that the observed lesions, indicating damage to nodes in these specific functional networks, may be associated with the observed effects on voluntary respiratory control (Supplementary Figure 6).

A complementary principal component analysis (PCA) revealed that the aforementioned four components explained 89.7% of the variance. When a fifth component was added, this value increased to 93.5%. However, the fifth component was at the boundary of the inclusion criteria. The PCA components appeared as mixtures of various networks (Supplementary Figure 7).

*PALM Analysis*

In addition to the network analysis, we also conducted a PALM analysis according to Joutsa et al. (2022) to investigate the correlations between lesion-related networks and behavior. This analysis was intended to provide further insights into the specific effects of the lesions. Unfortunately, the PALM analysis did not yield significant results, likely due to the small sample size. Despite the use of various statistical correction methods such as False Discovery Rate (FDR), Family-Wise Error (FWE), and Threshold-Free Cluster Enhancement (TFCE), only one test achieved a p-value approaching the threshold of 0.95 for statistical significance. Overall, the PALM analysis does not provide sufficient evidence to draw firm conclusions about the relationships between lesion networks and behavioral performance.

In summary, the Functional Network Analysis provided insights into the role of specific cortical networks in voluntary respiratory control. Although the PALM analysis did not allow for definitive conclusions, the network mapping and PCA contributed significantly to understanding the complex relationships between brain lesions and network functions.

**Supplementary Table 1.**

*Baseline and Task-Related Physiological Responses*

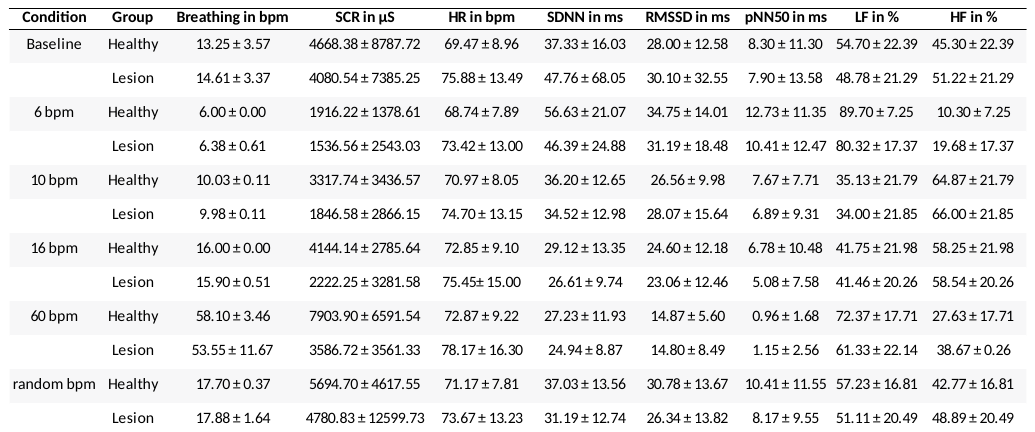

*Note.* The table presents mean ± SD for performed breathing, SCR and HRV for both healthy and lesion participants. SCR and HRV refer to each 2-minute episode. Breathing in bpm refers to breaths within a one-minute repetitive cycle; SCR in µS = skin conductance response in microsiemens; HR in bpm = heart rate in beats per minute; SDNN in ms = standard deviation of NN intervals in milliseconds; RMSSD in ms = root mean square of successive differences in NN intervals; pNN50 in ms = percentage of successive NN intervals that differ by more than 50 milliseconds; LF in % = low frequency power in percentage; HF in % = high frequency power in percentage.

**Supplementary Table 2.**

*Multilevel Task Correlation Matrices for Each Group Based on Spearman's Rho*

**Breathing Task**

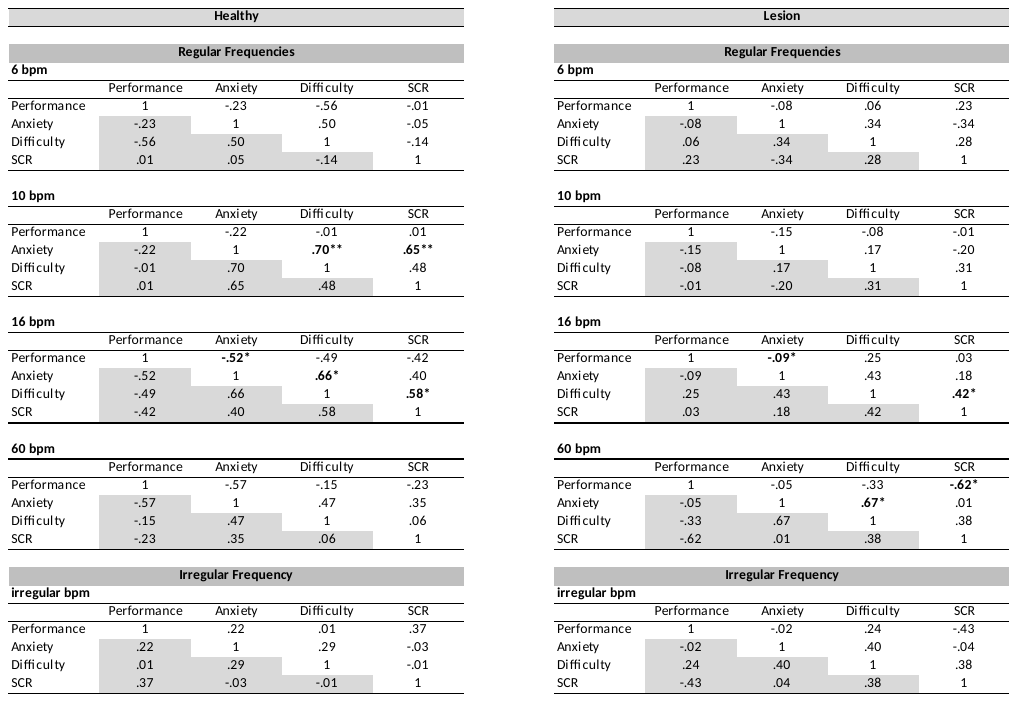

**Motor Task**

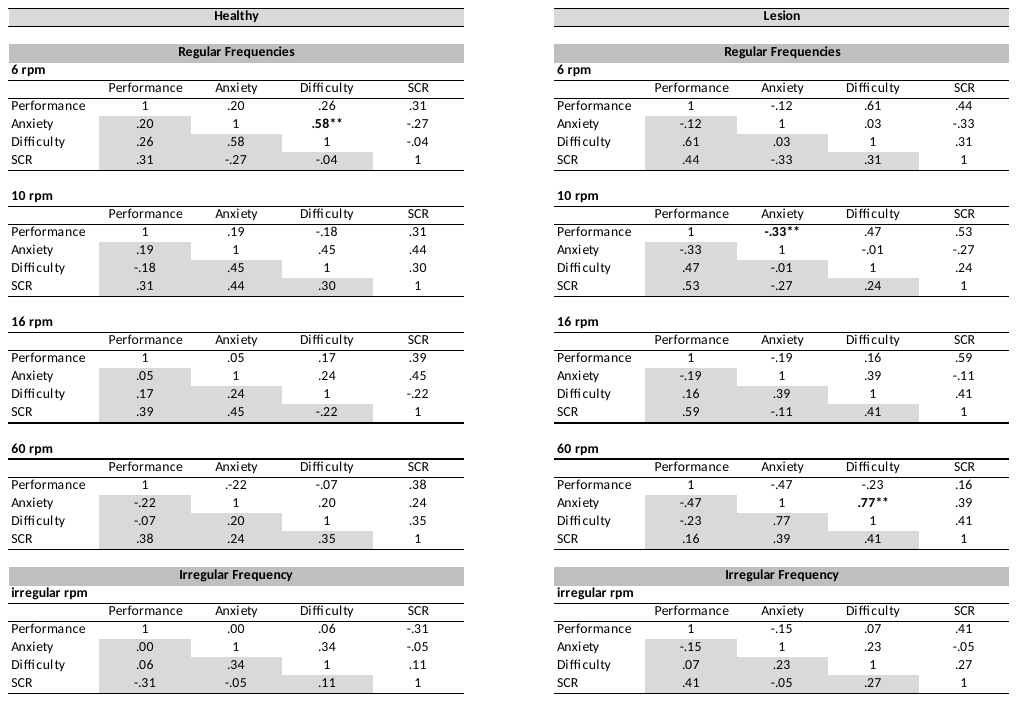

*Note.* Table presents Spearman's Rho correlation coefficients between the following variables: Performance (i.e., cross-correlation between instructed and recorded signal), Anxiety, Difficulty, and Skin Conductance Response for the healthy and lesion participant groups under the Breathing and Motor Task conditions. Asterisks denote the level of significance (* p<.05, ** p<.01, *** p<.001). Bpm = breaths per minute; Rpm = rotations per minute; SCR = Skin Conductance Rate.

**Supplementary Table 3.**

*Between Group HRV Parameters in Time- and Frequency Domain Across Regular Tasks*

| Variables | Lesion (n = 20) | Healthy (n = 20) | t-ratio *(df)* | p-value |
| --- | --- | --- | --- | --- |
|  | Mean ± SD | Mean ± SD |  |  |
| Respiration Task |  |  |  |  |
| AHR (bpm) | 75.4 ± 14.26 | 71.4 ± 8.59 | 2.19 *(158)* | .030* |
| RMSSD (ms) | 24.28 ± 15.31 | 25.20 ± 12.85 | -0.41 (*158*) | .683 |
| SDNN (ms) | 33.12 ± 17.43 | 37.05 ± 18.71 | -1.38(*158)* | .171 |
| pNN50 (%) | 5.88 ± 9.21 | 7.03 ± 9.49 | -0.78 *(158)* | .434 |
| pHF (%) | 45.72 ± 27.09 | 40.26 ± 28.66 | 1.24 *(158)* | .218 |
| pLF (%) | 54.28 ± 27.09 | 59.74 ± 28.66 | -1.24 *(158)* | .218 |
| Hand Motor Task | Lesion (n = 20) | Healthy (n = 18) |  |  |
| AHR (bpm) | 74.12 ± 10.40 | 71.85 ± 8.36 | 1.51 *(154)* | .134 |
| RMSSD (ms) | 23.33 ± 14.42 | 22.76 ± 12.19 | 0.26 *(154)* | .793 |
| SDNN (ms) | 27.57 ± 11.27 | 30.66 ± 16.00 | -1.40 *(154)* | .163 |
| pNN50 (%) | 6.68 ± 9.31 | 5.75 ± 7.91 | 0.67 *(154)* | .503 |
| pHF (%) | 52.01 ± 23.05 | 44.42 ± 19.80 | 2.28 *(154)* | .024* |
| pLF (%) | 47.99 ± 23.05 | 55.58 ± 19.80 | -2.28 *(154)* | .024* |

*Between Group HRV Parameters in Time- and Frequency Domain Across Irregular Tasks*

| Variables | Lesion (n = 20) | Healthy (n = 20) | t-ratio *(df)* | p-value |
| --- | --- | --- | --- | --- |
|  | Mean ± SD | Mean ± SD |  |  |
| Respiration Task |  |  |  |  |
| AHR (bpm) | 73.67 ± 13.23 | 71.17 ± 7.81 | 0.47 *(37)* | .472 |
| RMSSD (ms) | 26.34 ± 13.82 | 30.78 ± 13.67 | -1.02 (38) | .314 |
| SDNN (ms) | 31.19 ± 12.74 | 37.03 ± 13.56 | -1.40 (38*)* | .169 |
| pNN50 (%) | 8.17 ± 9.55 | 10.41 ± 11.55 | -0.67 *(37)* | .509 |
| pHF (%) | 48.89 ± 20.49 | 42.77 ± 16.81 | 1.03 *(38)* | .301 |
| pLF (%) | 51.11 ± 20.49 | 57.23 ± 16.81 | -1.03 *(37)* | .309 |
| Hand Motor Task | Lesion (n = 18) | Healthy (n = 19) |  |  |
| AHR (bpm) | 73.58 ± 10.08 | 71.78 ± 8.74 | 0.58 *(34)* | .567 |
| RMSSD (ms) | 39.80 ± 31.48 | 34.73 ± 42.63 | 0.41 *(35)* | .684 |
| SDNN (ms) | 38.91 ± 29.39 | 37.09 ± 30.41 | 0.19 *(35)* | .854 |
| pNN50 (%) | 7.89 ± 11.78 | 5.67 ± 11.52 | 0.57 *(35)* | .578 |
| pHF (%) | 55.87 ± 21.48 | 47.91 ± 23.67 | 1.01 *(34)* | .319 |
| pLF (%) | 44.13 ± 21.48 | 52.10 ± 23.67 | -1.01 *(34)* | .319 |
| *Note.* AHR: Average heart rate; bpm: Beats per minute; df: Degrees of freedom; RMSSD: Root mean sum of squared distance; SDNN: Standard deviation of the NN interval; pNN50: Proportion of consecutive RR intervals that differ by more than 50ms; pLF: Low-frequency power; pHF; High-frequency power; SD: Standard deviation. | | | | |

**Supplementary Figure 1.**

*Respiratory and Hand Motor Regulation Performance for Each Trial Condition on the Regular Task.*
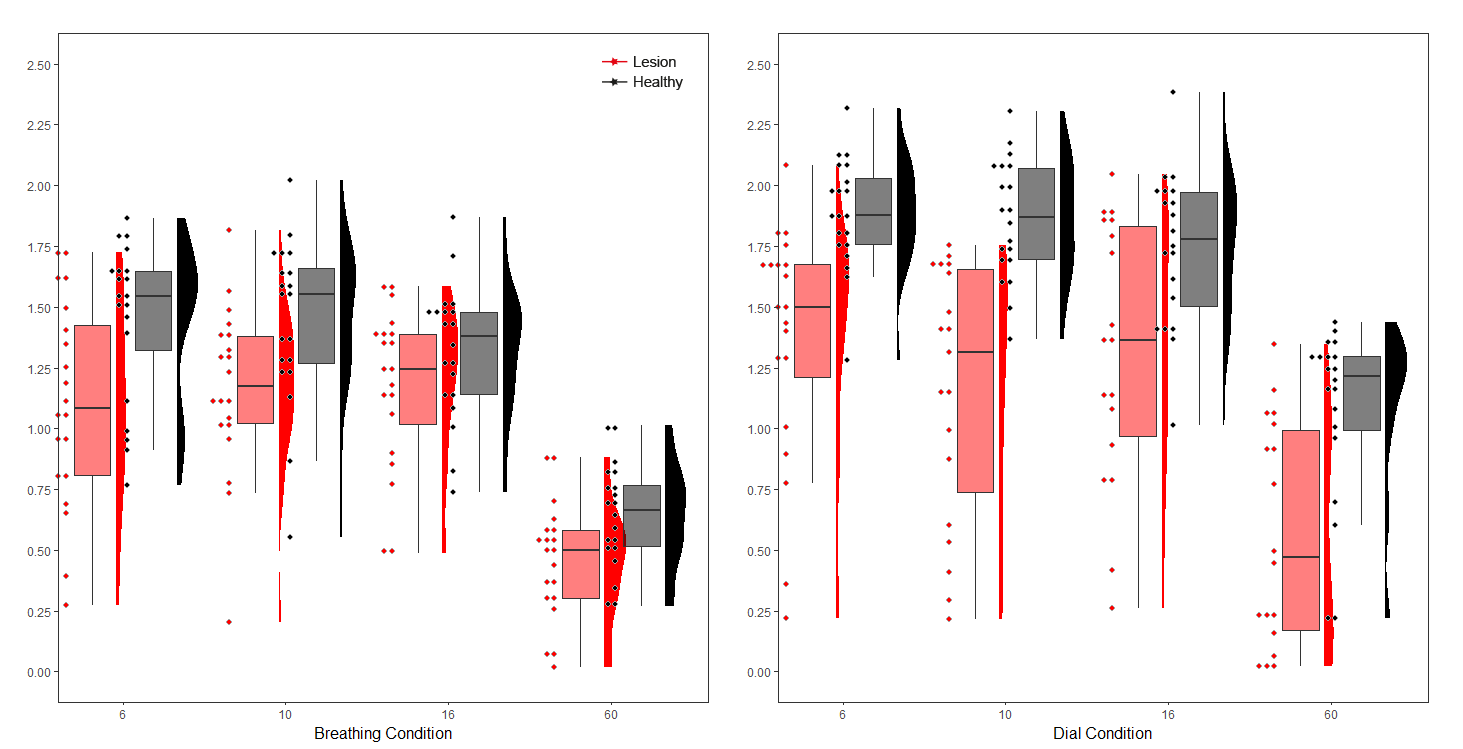

Fisher`s Z-transformed max cross correlation values

Fisher`s Z-transformed max cross correlation values

*Note.* Regulatory performance is represented on the Y-axis by Fisher`s Z-transformed max cross correlation values (range: 0 to 2.65).

**Supplementary Figure 2.**

*Analysis of Task Difficulty Ratings during Respiration and Motor Tasks Between Healthy Individuals and Individuals with Lesions*

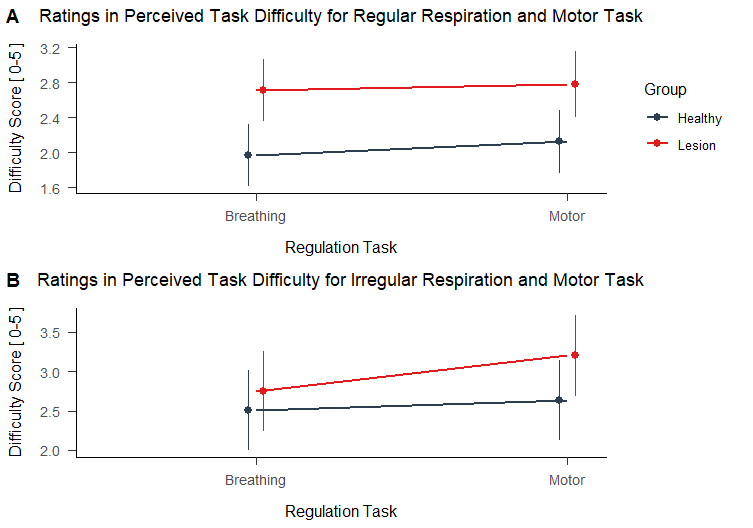

p = 0.004

*Note*. See manuscript for a detailed description of the statistical analysis results.

**Supplementary Figure 3.**

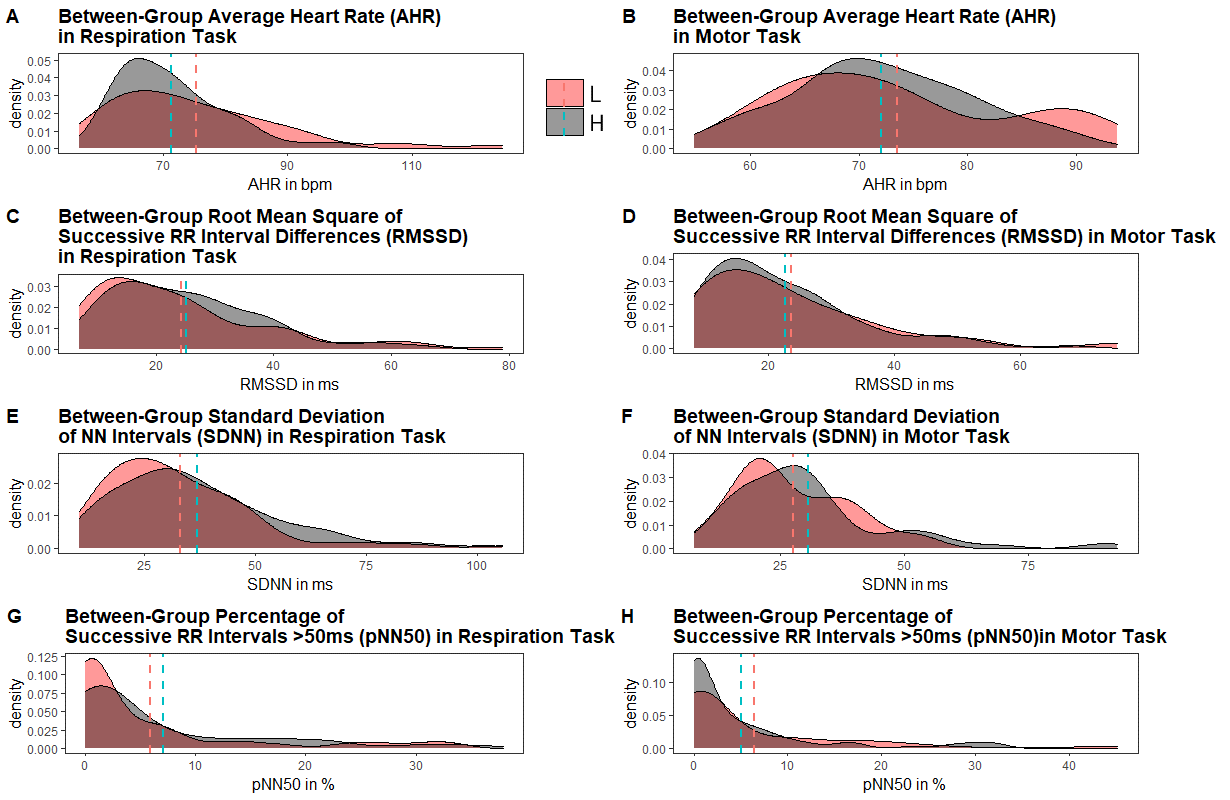
*Density for HRV Time Domain Parameters Between Experimental Groups for Regular Respiratory and Motor Regulation Tasks*

*Note*. The left column shows the time domain HRV parameters (A) AHR, (C) RMSSD, (E) SDDN and (G) pNN50 for the regular respiratory regulation tasks, while the right column shows the time domain parameters (B) AHR, (D) RMSSD, (F) SDDN and (H) pNN50 for the regular motor control tasks; dashed lines correspond to group mean; L: Lesion; H: Healthy.

**Supplementary Figure 4.**

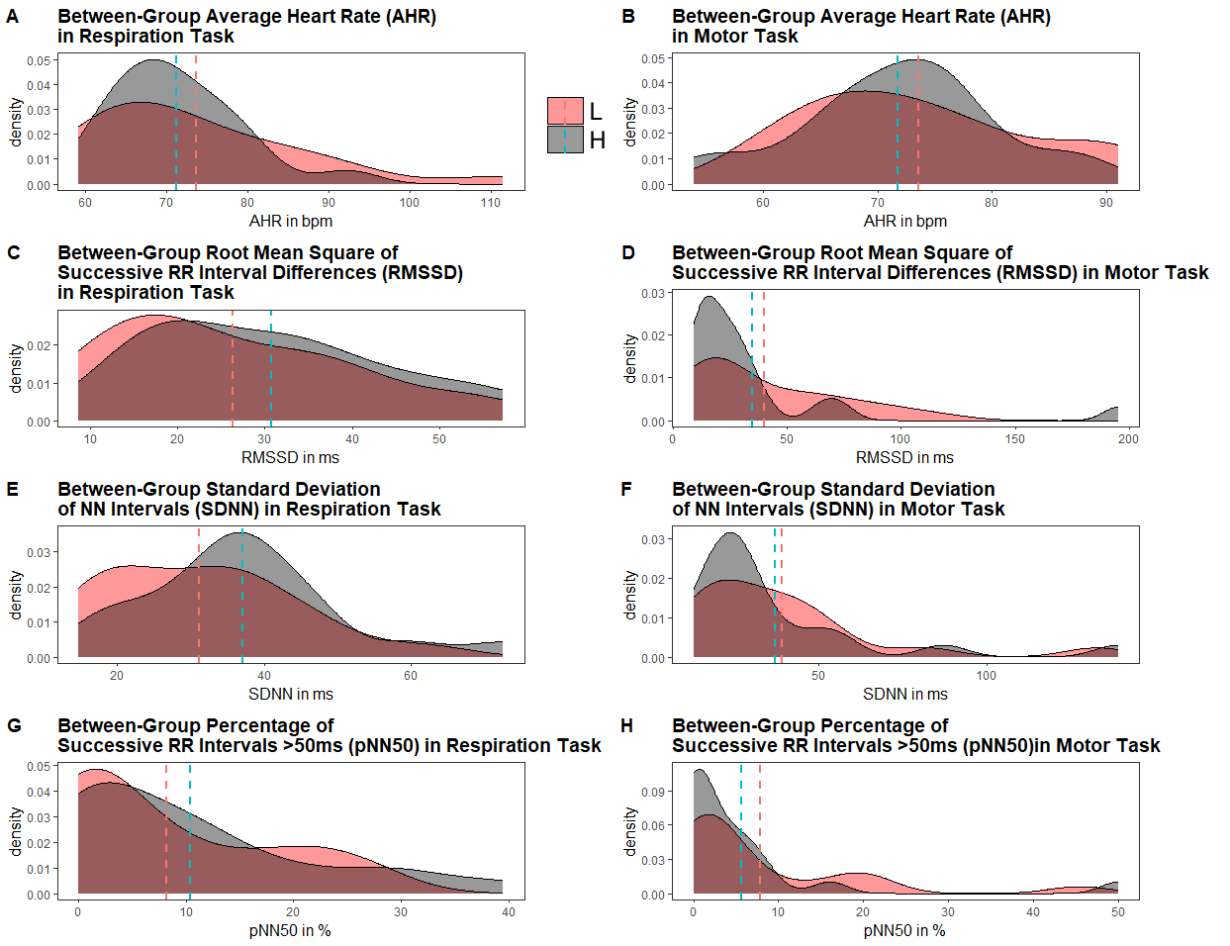
*Density for HRV Time Domain Parameters Between Experimental Groups for Irregular Respiratory and Motor Regulation Task*

*Note*. The left column shows the time domain HRV parameters (A) AHR, (C) RMSSD, (E) SDDN and (G) pNN50 for the irregular respiratory regulation task, while the right column shows the time domain parameters (B) AHR, (D) RMSSD, (F) SDDN and (H) pNN50 for the irregular motor control task; dashed lines correspond to group mean; L: Lesion; H: Healthy.

**Supplementary Figure 5.**

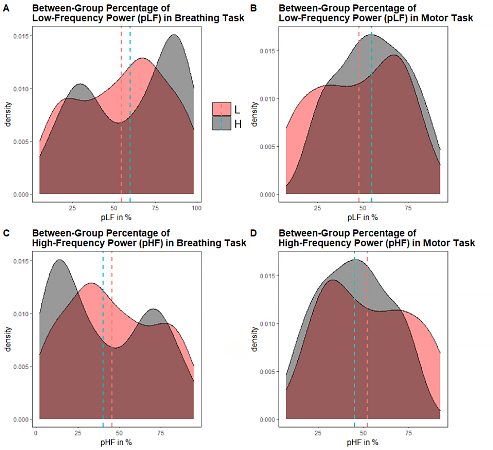
*Density for HRV Frequency Domain Parameters Between Experimental Groups for Regular Respiratory and Motor Regulation Tasks*

*Note*. The left column shows the frequency domain HRV parameters (A) pLF and (C) pHF, while the right column shows the time domain parameters (B) pLF and (D) pHF for the regular motor control task; dashed lines correspond to group mean; L: Lesion; H: Healthy.

**Supplementary Figure 6.**

*Density for HRV Frequency Domain Parameters Between Experimental Groups for Irregular Respiratory and Motor Regulation Task*

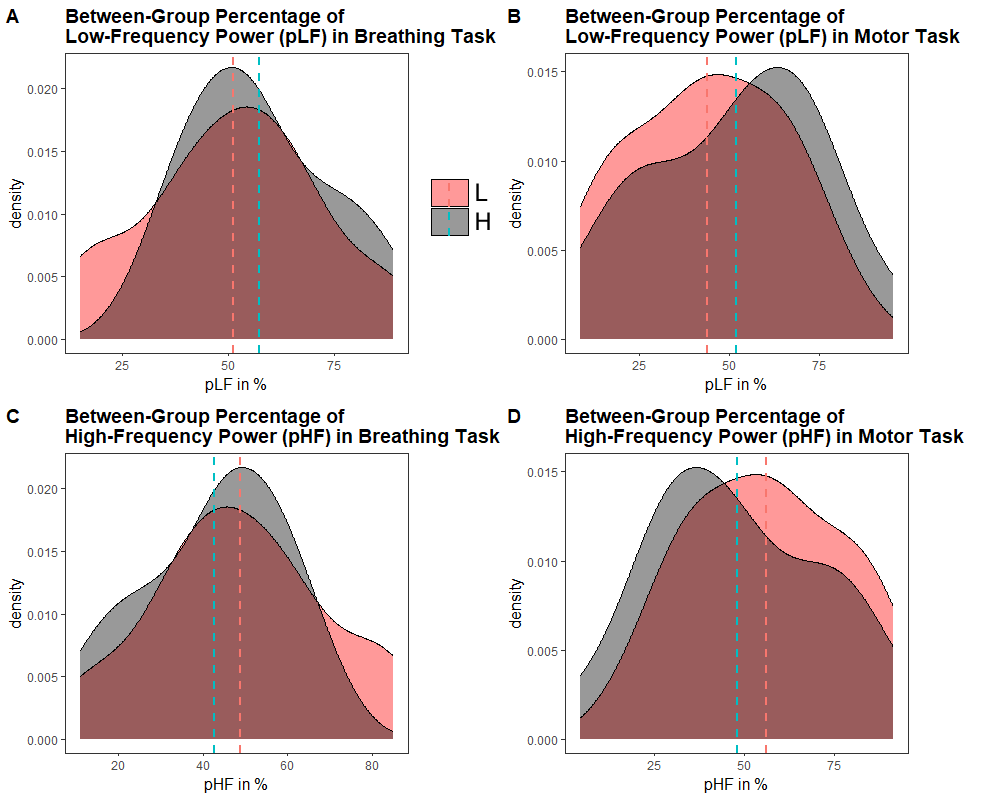

*Note*. The left column shows the frequency domain HRV parameters (A) pLF and (C) pHF, while the right column shows the time domain parameters (B) pLF and (D) pHF for the irregular motor control task; dashed lines correspond to group mean; L: Lesion; H: Healthy.

**Supplementary Figure 7.**

*Spatial Correlation Patterns Between Brain Lesions and Yeo7 Network Domains*
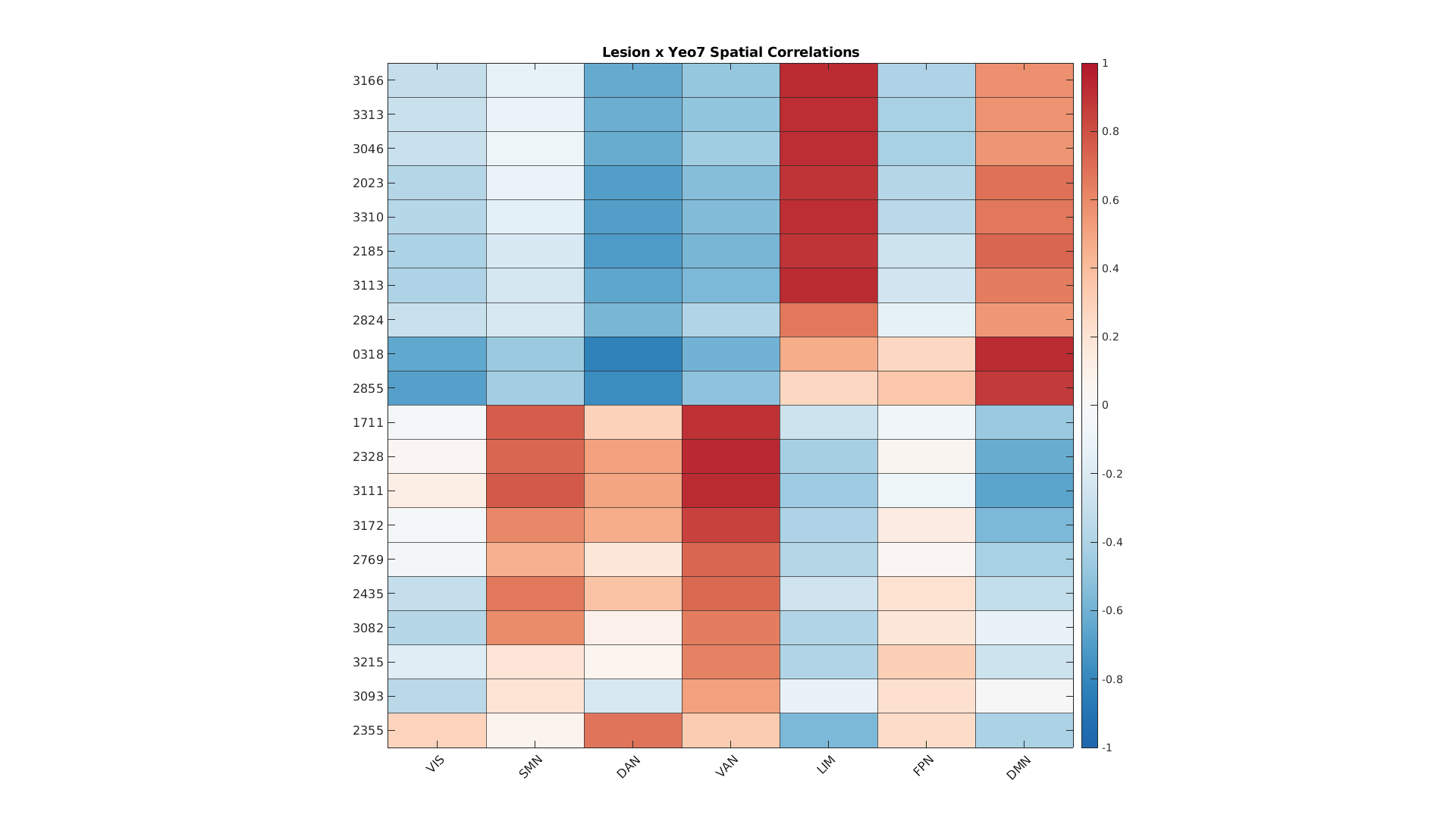

*Note.* The heatmap illustrates the spatial correlation patterns between individual brain lesions of participants (listed vertically) and the seven Yeo7 network domains (listed horizontally). The color scale indicates the correlation range from -1 (dark blue) for a strong negative correlation, to +1 (dark red) for a strong positive correlation. Intensities of color denote the degree of correlation, with dark red signifying high positive associations between lesion locations and networks, and dark blue indicating strong negative associations. Notably, significant positive correlations were observed in the Default Mode Network (DMN), Ventral Attention Network (VAN), Limbic Network (LIM), and the Somatomotor Network (SMN), suggesting their potential involvement in voluntary respiratory control. The correlation range from -1 to +1 highlights the variability and specificity of the interactions between network domains and brain lesions within this cohort*.*

**Supplementary Figure 8.**

Scree Plot of the Principal Component Analysis (PCA) for Variance Explained in Cortical Networks of Voluntary Respiratory Control

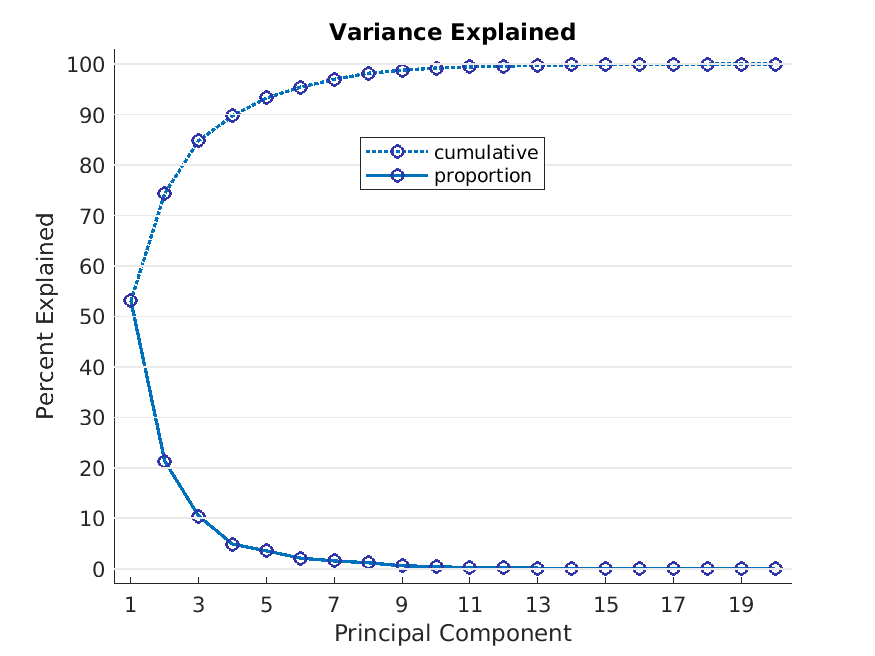

*Note.* The scree plot illustrates the percentage of variance explained by each principal component in the principal component analysis (PCA). The solid line indicates the proportion of variance explained by each individual component, while the dotted line shows the cumulative variance explained. The first four components account for 89.7% of the variance, with a marginal increase to 93.5% when a fifth component is included, albeit at the threshold of inclusion criteria.

**Supplementary Figure 9.**

Spatial visualization of the distribution of principal components 1 through 5.

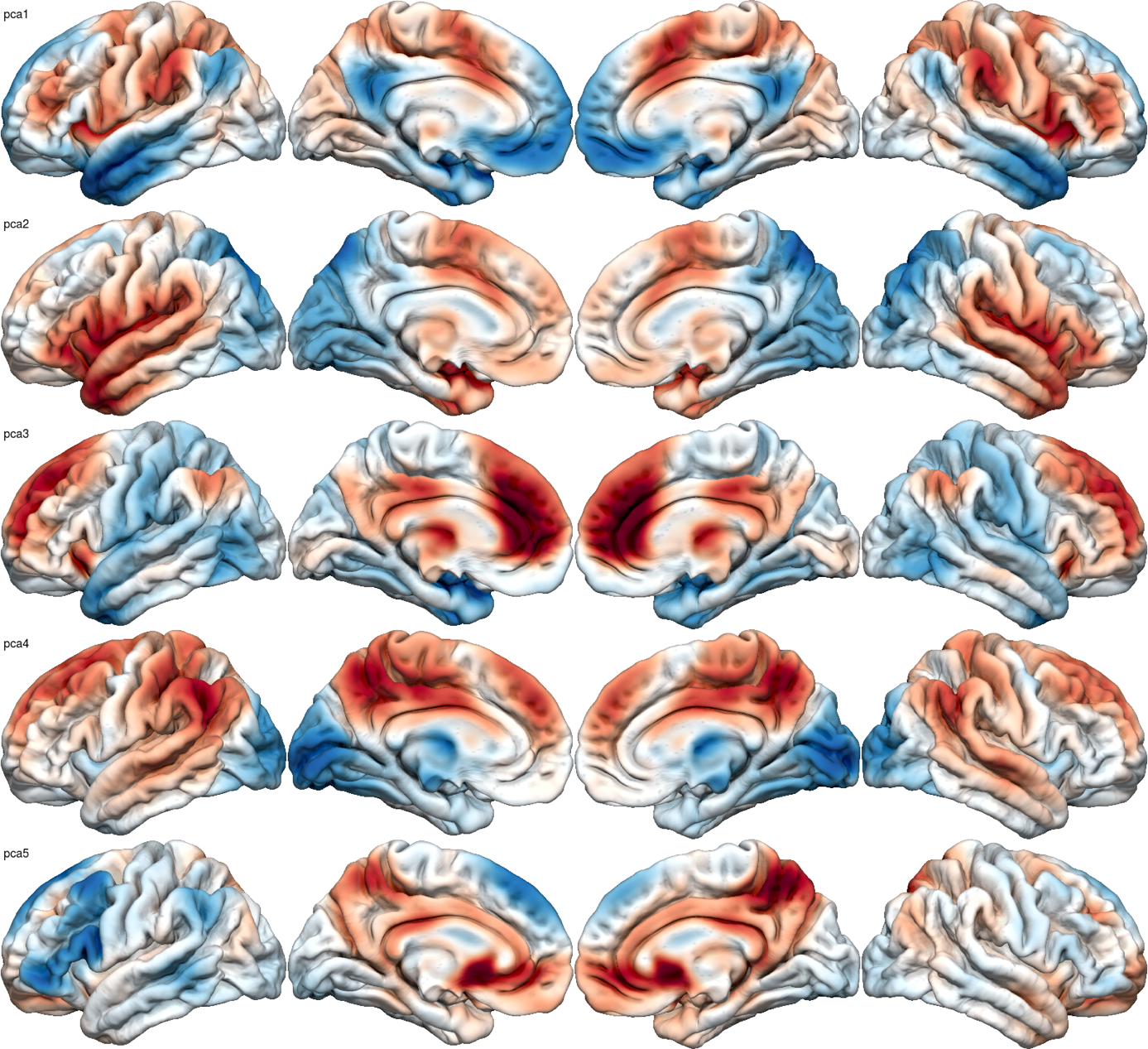

*Note. Variance in functional connectivity attributable to lesion location across the five principal components tended to be grouped in approximately four of the Yeo7 networks, namely the Limbic Network (LN), Default Mode Network (DMN), Somatomotor Network (SMN) and Ventral Attention Network (VAN).*

Variability. In 2019 Computing in Cardiology Conference (CinC). *Computing in*

*Cardiology*. Retrieved from https://doi.org/10.22489%2Fcinc.2019.032
